# Acute exposure of mouse and human hippocampal slices to tau oligomers reversibly impairs sharp-wave ripples

**DOI:** 10.64898/2026.04.05.716545

**Authors:** Chrystalleni Vassiliou, Janine Hochmair, Rithika Sankar, Ana Carolina Odebrecht Vergne de Abreu, Julia Onken, Thomas Sauvigny, Pawel Fidzinski, Susanne Wegmann, Camin Dean

**Author notes:** **Correspondence:** Camin Dean, Hufelandweg 14, Ebene 1, Campus Charité Mitte, 10117 Berlin.

## Abstract

Sharp-wave ripple (SWR) oscillations are crucial for memory consolidation and deteriorate in Alzheimer’s disease (AD). Tau oligomers are suggested to lead to synaptic and neuronal degeneration in AD, but their effects on SWRs are unknown. To study this, we prepared mouse and human hippocampal slices and bath-applied tau oligomer preparations after spontaneous SWR generation. In human slices, acute exposure to tau resulted in decreased ripple duration, whereas in mouse slices it was SWR rate, amplitude, and power that decreased, sparing duration. In a different set of experiments, mouse slices were pre-incubated directly in either tau-ACSF or control-ACSF right after slicing for 2.5–5.5 hours, resulting only in diminished SWR rate. These effects were specific to the presence of β-sheets, as a different tau preparation that lacked β-sheets failed to alter SWRs. This method is therefore suitable to study SWR alterations after short-term exposure to different tau and/or Aβ species, allows a higher throughput screening of possible therapeutics compared to in vivo animal experiments, and permits direct comparison of SWR alterations in mice and humans.

## Introduction

Sharp-wave ripples (SWRs) are hippocampal oscillations essential for memory consolidation. They are observed during non-rapid eye movement (NREM) sleep and awake resting periods. They consist of two co-occurring oscillations: sharp-waves (SWs), which are large-amplitude non-periodic deflections, and ripples, 70–250 Hz oscillations in humans and 100– 250 Hz in mice (1). During SWRs, neuronal firing patterns representing a previous experience are reactivated (2), leading to strengthening of memory-specific connections between the firing neurons, i.e., long-term potentiation (LTP; (3)). Simultaneously, SWRs trigger widespread long-term synaptic depression (LTD) of all other connections, to rescale synaptic weights, which is necessary for subsequent learning (4,5). Importantly, inhibition of SWRs impairs memory (6) while their enhancement improves memory (7,8).

Alzheimer’s disease (AD), characterized by intracellular tau-containing neurofibrillary tangles and extracellular deposition of amyloid plaques, is the most prevalent type of dementia and is associated with SWR impairments (9). For example, a transgenic mouse model that overexpresses a mutant form of tau (rTg4510) shows decreased ripple amplitude and occurrence rate but spares ripple duration (10,11). Tau oligomers, occurring prior and in addition to overt tau aggregating in neurofibrillary tangles, are synaptotoxic and may spread tau pathology across neuronal networks (12–14). Subcortical injection of tau oligomers, but not monomers or fibrils, leads to neuronal degeneration and memory impairment in wild-type mice (15), suggesting that the tau oligomers are the toxic species. In agreement, LTP magnitude decreases after exogenous application of tau oligomers in mouse hippocampal slices (16,17). However, to our knowledge, it remains unknown whether short-term exposure to tau oligomers causes acute SWR dysregulation.

In this study, we bath-applied human tau oligomer preparations on hippocampal slices from humans and mice, hypothesizing that even a 30-minute exposure to tau would be enough to impair SWRs. We found that even though SWRs were diminished in both species, it was ripple duration that was decreased in human slices but SWR rate of occurrence, amplitude, and power in mouse slices. This method permits investigation of early tau-induced network dysfunction, which is not possible in constitutive transgenic animal models, in an ex vivo set-up of the isolated hippocampus. This procedure could therefore be utilized to test SWR changes caused by specific tau and/or Aβ species, as well as to screen possible therapeutic agents.

## Methods

### Tau protein purification

Recombinant human full-length tau (2N4R isoform) was expressed and purified following established protocols (18,19): Protein expression in E. coli BL21 Star (DE3; Invitrogen) was induced with 0.5 mM IPTG at OD_600_=0.6 for ∼3 h at 37°C. Cells were harvested, resuspended in lysis buffer (20 mM MES, 1 mM EGTA, 0.2 mM MgCl_2_, 1 mM PMSF, 5 mM dithiothreitol (DTT), protease inhibitors (Pierce Protease Inhibitor Mini Tablets, EDTA-free)) and lysed using a French press. After adding 500 mM NaCl and boiling at 95°C for 20 min, cell debris was removed by centrifugation and the supernatant was dialyzed against low salt buffer (20 mM MES, 50 mM NaCl, 1 mM MgCl_2_, 1 mM EGTA, 2 mM DTT, 0.1 mM PMSF, pH 6.8), filtered (using a 0.22 μm membrane filter), run through a cation exchange column (HiTrap SP HP, 5 ml, GE Healthcare), and eluted with a high salt buffer (20 mM MES, 1000 mM NaCl, 1 mM MgCl_2_, 1 mM EGTA, 2 mM DTT, 0.1 mM PMSF, pH 6.8). Fractions containing tau were pooled, concentrated using spin column concentrators (Pierce Protein concentrators; 10–30 kDa MWCO, Thermo Fisher Scientific), and run through a size exclusion column (Superose 6 10/300, GE Healthcare). Fractions containing purified monomeric tau were concentrated as before, buffer exchanged to PBS containing 1 mM DTT, aliquoted, flash frozen, and stored at −80°C.

### Preparation of human tau oligomers

Two different methods were used to prepare tau oligomers, one using a standard preparation with heparin (19) and one using hydrogen peroxide (H_2_O_2_ (16)). The resulting tau oligomers were assessed using a Thioflavin-T aggregation assay.

#### Oligomeric tau prepared with heparin (otau)

60 μM of tau monomers were mixed with 0.7 mg/ml heparin (Applichem; MW= 8–25 kDa) in PBS with 2 mM DTT. The sample was then incubated for 6 days at 37°C to allow full aggregation, with fresh 2 mM DTT added every second day. Dialysis against sterile PBS (Pur-A-Lyzer, 10 kDa MWKO, Sigma-Aldrich) overnight at 4°C was followed by sonication for 30 min to break tau aggregates into oligomers. The sample was then immediately aliquoted and flash frozen in liquid nitrogen, to prevent further tau aggregation, and stored at -80°C. For the batch used to pre-incubate mouse slices with tau and in human tissue experiments, instead of buffer exchange, the samples were ultracentrifuged to remove tau monomers before resuspending in sterile PBS and sonicating to break tau aggregates into oligomers.

#### Oligomeric tau prepared with H_2_O_2_ (otau-H_2_O_2_)

DTT was removed from tau preparations by dialyzing sterile PBS twice (first for 2 h, then overnight) at 4°C using a dialysis device (Pur-A-Lyzer, Sigma-Aldrich) with a 12⍰kDa molecular weight cut-off (MWCO). If needed (to reach a concentration of 50 µM needed for the aggregation reaction) monomers were further concentrated using 3 kDa MWCO concentrators (Pierce protein concentrators, Thermo Fisher Scientific) at 10,000 x g at 4°C. Afterwards, 183 µM tau monomers were incubated with 1 mM H_2_O_2_, 1 x Halt Protease inhibitor mix (cat. #78429, Thermo Fisher Scientific) and 0.01% NaN_3_ at 25°C in a Thermoblock for 20 h at 250 rpm. The sample was then centrifuged for 20 min at 10,000 x g, and the supernatant buffer exchanged against PBS using dialysis as before. The sample containing tau oligomers and monomers was sonicated for 15 min in a water bath. The resulting tau oligomer preparation was aliquoted, flash frozen with liquid nitrogen, and stored at -80°C.

### Thioflavin-T aggregation endpoint assay

This assay was performed as described before (19). Briefly, to detect and quantify the presence of β-sheet containing tau oligomers, 4⍰μM of each tau oligomer preparation was incubated with 50⍰μM Thioflavin-T (ThioT, Sigma-Aldrich), a dye which upon binding β-sheets increases fluorescent intensity, in 384-well μClear plates (Greiner Bio-One) in triplicates. Thio-T fluorescence (excitation at 440⍰nm, emission at 485⍰nm) was recorded at 37°C using an Infinite M Plex plate reader (Tecan), following a 5-second shaking step to ensure sample mixing. Fluorescence was recorded 10 min after ThioT addition. Oligomeric tau prepared with heparin had a Thio-T fluorescence higher than controls (tau monomers or Thio-T buffer; *p* < 0.0001; Fig. 1), indicating aggregation and β-sheet formation. Thio-T fluorescence of the otau-H_2_O_2_ was similar to controls, indicating a lack of β-sheet formation.

**Figure 1:**
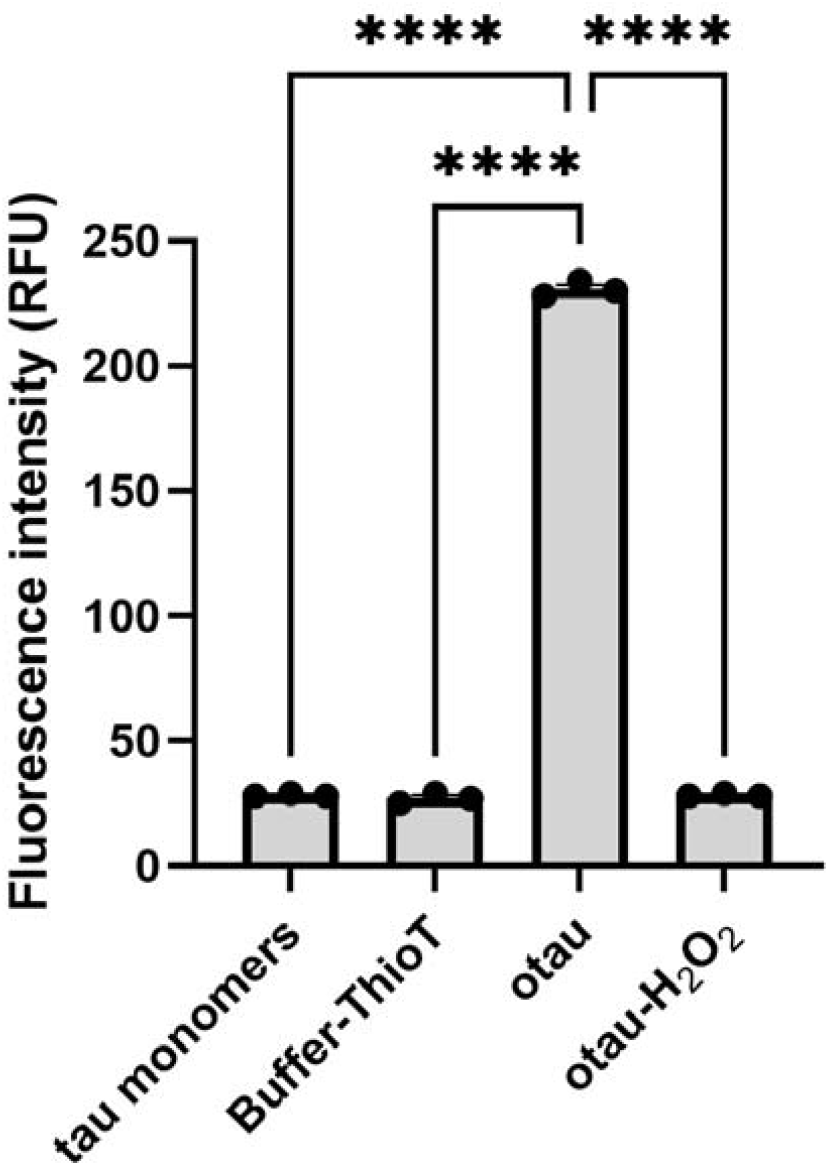
Tau β-sheets were present only in oligomeric tau prepared with heparin. Thioflavin-T fluorescence of oligomeric tau preparations was higher than tau monomers and buffer-Thio-T controls in otau prepared with heparin (*F*(3,8) = 8812, *p* < 0.0001), but not in otau-H_2_O_2_, suggesting that β-sheet containing tau oligomers were present only in the otau prepared with heparin. Significance was tested by one-way ANOVA. Pairwise differences were evaluated with Tukey’s multiple comparisons test with **** = *p* < 0.0001. N = 3 plates per preparation. Graph shows means ± standard errors (SEs).

### Subjects

#### Mice

Male and female C57BL6/J mice 8–14 weeks old were acquired either internally from Charité Forschungseinrichtungen für Experimentelle Medizin (FEM) or externally from Charles River Laboratories (strain code #632, RRID:SCR_003792). They were group housed whenever possible and kept on a 12-hour light-dark cycle with ad libitum access to food and water. More information on slices and mice used in each experiment can be found in Table 1. All procedures described here were performed in accordance with the German animal welfare laws (Tierschutzgesetz), the Directive 2010/63/EU of the European Parliament and of the Council of 22 September 2010 on the protection of animals used for scientific purposes, as well as the Ethics Committee of the Charité under the guidelines of the Berlin state authorities (T-CH 0013/20).

**Table 1.** Information on mouse slices per experiment.

| Experiment | Mouse age and sex | Slices |
| --- | --- | --- |
| Acute application of otau | 8 mice (4 male, 4 female), 8 weeks old | 16 slices (before, during, and after exposure to tau), 1–3 slices per mouse |
| Pre-incubation with or without otau | 8 mice (4 male, 4 female), 10–14 weeks old | 16 control, 18 tau, 1–4 slices per mouse per condition (pre-incubated with or without tau) |
| Acute application of otau-H<sub>2</sub>O<sub>2</sub> | 5 mice (3 male, 2 female), 8–9 weeks old | 18 slices (before, during, and after exposure to tau), 2–4 slices per mouse |
| Pre-incubation with or without otau-H<sub>2</sub>O<sub>2</sub> | 2 female mice, 10–11 weeks old | 5 control, 7 tau, 2–4 slices per mouse per condition (pre-incubated with or without tau) |

#### Human patients

Human hippocampal tissue was received from four male 7–75 year old patients undergoing hippocampectomy for the treatment of drug-resistant epilepsy or tumor removal. Written consent allowing the scientific use of the resected tissue, which would otherwise be discarded, was provided by all patients. Table 2 lists patient age, reason for hippocampectomy, and number of slices used in this study. All procedures were approved by the ethics committees of the Charité – Universitätsmedizin Berlin and the medical association of Hamburg for University Medical Center Hamburg-Eppendorf (EA2/111/14, EA2/064/22 in reference to EA4/206/20; 2023-200674-BO-bet in reference to EA2/111/14, respectively).

**Table 2.** Information on human hippocampal slices.

| Patient sex and age | Patient condition | Surgery location | Tissue transportation time | Slices recorded |
| --- | --- | --- | --- | --- |
| Male, 19 years old | Mutation in the ACTB gene, including mild cognitive disorder, intractable epilepsy | Berlin | 30 minutes | 3 |
| Male, 75 years old | Low-grade epilepsy-associated tumor (cavernoma) | Berlin | 30 minutes | 4 |
| Male, 23 years old | Temporal lobe epilepsy with hippocampal sclerosis | Berlin | 30 minutes | 3 |
| Male, 7 years old | Temporal lobe epilepsy with dysembryoplastic neuroepithelial tumor (DNET) | Hamburg | 5 hours | 2 |

### In vitro electrophysiology

#### Slice preparation

##### Mouse tissue

Similar experimental procedures have been performed in previous studies (20,21). Mice were decapitated after isoflurane (Forene, cat. #B506, AbbVie) inhalation. The brains were extracted and placed on a cold plate with ice-cold (0.5–2°C) standard artificial cerebrospinal fluid solution (ACSF) containing, in mM: 119 NaCl, 2.5 KCl, 1 NaH_2_PO_4_, 1.3 MgSO_4_, 2.5 CaCl_2_, 26 NaHCO_3_, and 10 D-glucose. It was oxygenated by bubbling carbogen (95% O_2_, 5% CO_2_) at a pressure of 32 ± 1 mmHg, with pH under these conditions being 7.3–7.4. Both hippocampi were excised from each brain and placed one at a time on a tissue chopper (cat. #51425, Stoelting) with the ventral surface perpendicular to the blade. Four slices 450 μm thick were taken from each hippocampus. Slices were then directly transferred to an interface chamber inside a Farraday cage, where they were left to incubate for at least two hours in 31 ± 1°C oxygenated ACSF before recording.

##### Human tissue

Resected tissue from patients was transferred from the operating room in Berlin (surgeries took place in Charité – Universitätsmedizin Berlin, diagnosis in Evangelischen Krankenhaus Königin Elisabeth Herzberge) or Hamburg (Universitätsklinikum Hamburg-Eppendorf) in ice-cold choline-chloride ACSF containing, in mM: 110 choline chloride, 11.6 (+)-sodium L-ascorbate, 7 MgCl_2_, 2.5 KCl, 1.25 NaH_2_PO_4_, 0.5 CaCl_2_, 3.1 sodium pyruvate, 26 NaHCO_3_, and 10 D-glucose. This ACSF was only used for transport and slicing of the human tissue, while the standard ACSF described above was used for everything else. Transfer of the tissue from the operating room to the vibratome took roughly 30 minutes for surgeries in Berlin and 5 hours for the surgery in Hamburg. Once the tissue was brought to Charité Campus Mitte, it was placed on a vibratome (model VT1200S, Leica Microsystems) with choline-chloride ACSF and 3–11 slices 450 μm thick were taken from each tissue sample. Slices were placed on a transfer chamber containing standard oxygenated ACSF at 30 ± 5°C in submerged conditions until all slices were collected and moved to a different building in the same campus. They were then transferred to the same interface chamber used for mouse slices and incubated for at least two hours in 31 ± 1°C oxygenated ACSF before recording 2–4 slices per sample.

##### Tau application

In all experiments, 25 nM tau oligomers in ACSF were bath-applied to hippocampal slices. Tau exposure started either right after slicing (pre-incubation with tau-ACSF) or after recording SWRs in tau-free ACSF (acute tau-ACSF application).

##### Pre-incubation with tau-ACSF

Hippocampal slicing was performed in tau-free ACSF. Slices were then immediately separated into two groups and incubated for at least two hours in two different interface chambers, one with tau-ACSF and the other with tau-free ACSF. Afterwards, slices with spontaneous SWRs from each condition were identified and SWRs were recorded for 35 minutes from each slice. Two minutes from the last 27–35 minutes of each slice recording were extracted for analysis. This was done to allow the tissue and SWRs to recover after the recording electrode entered the slice and select a time with stable baseline and as little artefacts (sharp negative deflections roughly 0.2 ms long that appear simultaneously on at least two recording channels) as possible. If more slices with SWRs were present, more recordings were taken, resulting in total incubation time between 2.5–5.5 hours. Because the amount of tau was limited and this experiment required long tau application, the ACSF (∼100 mL tau-ACSF and ∼100 mL tau-free ACSF) was reused.

##### Acute tau-ACSF application

Hippocampal slices were prepared and incubated in tau-free ACSF for at least two hours. A 35-minute-long baseline (pre-tau) SWR recording was taken from slices with spontaneous SWRs. The ACSF was then switched to tau-ACSF and a second 35-minute-long recording (tau) was obtained. Finally, slices were switched back to tau-free ACSF for another 35-minute-long recording, to see if the effects of short-term tau application could be reversed by washing (post-tau). Two minutes from the final 27–35 minutes of each condition (pre, tau, and post) were extracted for analysis. The two minutes selected had the least number of artefacts and the most stable baseline, while allowing for roughly 30 minutes of exposure to tau or washout.

##### SWR recordings

Slices were kept at 31 ± 1°C in standard ACSF oxygenated with carbogen at 32 ± 1 mmHg. The ACSF was flowing at a rate of 1.3–1.5 ml/min. Under these conditions, SWRs are generated spontaneously after two hours of incubation. The CA1 pyramidal layer, where SW polarity is positive and ripple amplitude is the largest, was identified using a light microscope (model SZ61; Olympus Corporation). Local field potential (LFP) recordings were performed using platinum/iridium microelectrodes (0.1 MΩ, cat. #PI2PT30.1H5, except for acute otau application in mouse tissue where four slices were recorded with #PI20030.1A3 MicroProbes for Life Science). The LFP from the surface of the slice was examined, until the location in the CA1 pyramidal layer with the largest SW amplitude was identified. The electrode was then inserted roughly 200–250 μm deep into the slice. A differential AC amplifier (cat. #1700, A-M Systems) was used to capture frequencies between 0.1–5,000 Hz and amplify the signal 10,000-fold. The signal was digitized at 10 kHz with a Micro3 1401 data acquisition interface (Cambridge Electronic Design) and recorded using Spike2 software, version 8.11 (Cambridge Electronic Design).

##### Analysis of SWR properties

SPIKE 2v8.11 was used to extract two minutes per recording for analysis (as described above), which was performed in MATLAB 2023b (The MathWorks, Inc.) using a custom written code (https://github.com/ChrystalleniCy/SWR-analysis---Tau.git). For SWs, the signal was filtered between 1–25 Hz, while for ripples between 100–250 Hz for mouse slices and 70–250 Hz for human slices. Filtering was performed using the eegfilt.m script from the EEGLAB toolbox (22). The FIR filter coefficients had a length of 20,000 points for SWs and 7,000 for ripples. To avoid phase distortion, the filter was applied in both forward and reverse directions using MATLAB’s filtfilt function, resulting in a zero-phase filter.

While it is well established that rodent and primate ripples have different frequencies (1,23,24), other SWR parameters such as power (25) and rate (26,27) have also been suggested to differ between species. Therefore, the same SWR detection thresholds might not work well in both mouse and human slices. In order to check and account for that, we combined all pre-tau/control mouse recordings and compared them with the human pre-tau recordings. To estimate signal-to-noise ratio (SNR) in the two species before event detection takes place, a distribution-based metric of the entire two-minute SW- or ripple-filtered recording was used. The standard deviation (SD) of the distribution of amplitudes of every single point of a recording measures the spread of the entire distribution. It is therefore sensitive to the presence of high amplitude SW or ripple events and can be used to reflect the signal. On the other hand, the interquartile range (IQR) of this distribution captures the spread of 50% of the data and is not affected by the relatively rare events, so it can be thought to represent baseline noise. Hence, the SD/IQR ratio was used as an estimation of the signal-to-noise ratio (SNR) of each recording. The SNR was compared between mouse and human recordings using a generalized linear mixed effects model (glme; using the fitglme MATLAB function) with equation:

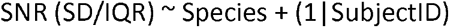

where (1|SubjectID) accounts for slices coming from the same subjects. While no difference was found in the SNR of the SW-filtered channel between mouse and human recordings, ripple SNR was significantly smaller in mice (0.86 ± 0.030; estimated marginal mean (EMM) ± standard error (SE)) compared to humans (1.22 ± 0.065; *p* = 4.1 × 10^-6^; Fig. 1A), with a ratio between human and mouse SNR = 1.42. Therefore, the same amplitude threshold was used for SW detection in both species (4.5 × SD), but a different threshold was used for ripple detection (for mice: 3 × SD, for humans: 3 × 1.42 = 4.3 × SD) and for the maximum ripple peak criterion (the amplitude each ripple event’s maximum peak had to surpass to be included in analysis, see below for details on ripple detection; for mice: 4 x SD, for humans: 4 x 1.42 = 5.7 x SD, Fig. 1B). Repeating this analysis using mouse data only from pre-tau recordings from acute experiments (where the ACSF was not reused) produced similar results (for SWs: *F*(1,44) = 0.78, *p* = 0.38; for ripples: *F*(1,44) = 19, *p* = 8.7 × 10^-5^, human = 1.22 ± 0.076, mouse = 0.84 ± 0.045, ratio = 1.46).

For detection of SWs, the baseline was normalized to zero by subtracting from all points the mode of the amplitude distribution of the SW-filtered signal. Subset A was defined as all points of the SW-filtered signal with amplitude -4 x SD_SW_signal to 0 μV, representing an underestimation of baseline noise which however lacks any part of the positive SW events. We took the SD of this subset (SD_A) and defined baseline noise (B) as all points of the SW-filtered channel within ± 4 x SD_A. Finally, candidate SW events were identified as peaks exceeding 4.5 x SD_B (SD of the baseline noise subset). The full-width at half-maximum (FWHM) of each candidate SW was calculated using an adapted version of fwhm.m (28), and only events with FWHM between 15–100 ms were considered for further analysis to exclude possible artefacts. The closest minimum point before and after each SW peak that was either the last minimum before another SW peak, or had an amplitude of less than 2 x SD_B was taken as the start and end of that SW peak respectively. A maximum of 3 minima before and 5 minima after each peak were checked. If none of these minima satisfied the aforementioned conditions, then the closest minimum to the peak was selected. If a SW at the start or end of the 2-minute recording snippet was incomplete, the start or end of the snippet were taken as that SW’s start or end, respectively. SW peak, start, and end detection was confirmed by manual inspection.

Detection of candidate ripple events was similar. The mode of the amplitude distribution of the ripple-filtered signal was first taken as the baseline of the recording, and points between mode ± SD_ripple_signal were taken as baseline noise of the ripple channel. The ripple channel was then rectified, making all peaks positive, and candidate ripple peaks were defined as peaks exceeding 3 x SD_baseline_noise for mouse recordings, or 4.3 x SD_baseline_noise for human recordings. To determine the maximum peak-to-peak time difference to group ripple peaks into a ripple event, the minimum ripple frequency was taken (100 Hz for mouse slices and 70 Hz for human slices), making the maximum ripple period 10 ms for mice and 14.3 ms for humans. Since each cycle has 2 peaks in the rectified channel, the maximum peak-to peak time is 5 ms for mice and 7.2 ms for humans. In order to allow for some missing peaks, this number was multiplied by 1.5, so ripple peaks closer than 7.5 ms for mouse slices and 11 ms for human slices were grouped into the same ripple event. Events with less than 6 peaks (= 3 cycles) were rejected. The maximum ripple peak of each ripple event was also identified and events with maximum peak less than 4 x SD_baseline_noise for mouse slices or 5.7 x SD_baseline_noise for human slices were also excluded from further analysis. Ripple start time was defined as the time of the first peak of each ripple event, end time as the time of the last peak of each event, and middle time as halfway between the two. For analysis of ripple power and frequency, 160 ms windows from the ripple-filtered signal centered at the ripple middle time were taken. If an event was too close to the start or end of the 2-minute recording snippet, then 160 ms were taken from the start or end of the snippet, respectively. The wavelet.m function (29) was used for wavelet transformation of each ripple event window, with parameters: sampling interval (dt) = 1/sampling frequency, resolution (dj) = 0.01, start scale (s0) = 0.002, number of scales (j1) = 5/dj, ‘morlet’ mother wavelet and zero-padding on. To compute ripple power, the absolute value of the wave output was squared. If two consecutive events had the same peak instantaneous ripple power and frequency (i.e., if two events were in the same 160 ms window, the stronger ripple would be counted twice) only the event whose time of peak instantaneous power was closer to the time of maximum ripple peak amplitude was included in analysis of ripple power and frequency (but both were included in analysis of other parameters), hence removing duplicates. Additionally, to avoid edge artefacts, events with time of peak instantaneous power within 20 ms of the edges of the 160 ms window were also excluded from analysis of ripple power and frequency only. The power was then averaged for each frequency across the entire ripple window duration. The maximum power of this time-averaged power spectrum was taken as the power of the candidate ripple event, and the frequency corresponding to this maximum power as the dominant frequency of the ripple event.

To extract SWR events for final analysis, the co-occurrence of SW and ripple events is necessary. Therefore, if the ripple middle time preceded the SW peak time and the SW start time preceded the ripple end time, or if the SW peak time preceded the ripple middle time and the ripple event start time preceded the SW end time (hence ripples and SWs overlapped in time), the SW and the ripple event were paired in a SWR event. Once a ripple event was paired to a SW event, analysis was moved to the next SW to prioritize pairing 1 ripple event with 1 SW event if two SWs were very close in time. Then, all SWs were tested for overlapping with remaining ripple events a second time. SWR events could therefore consist of 1 SW and 1 or 2 ripple events. Only SWs and ripples that were part of SWR complexes were analyzed further. Detection of candidate SW and ripple events, as well as analyzed SWR events, was confirmed visually.

For each slice, the SWR rate (SWR events per second), mean SW FWHM, mean SW amplitude, mean ripple amplitude (the amplitude of the maximum ripple peak per ripple event averaged over all ripple events), mean ripple power (peak time-averaged power of each ripple event averaged across events), mean frequency (dominant frequency of each ripple event averaged across events), mean ripple duration (ripple end time - ripple start time per ripple event), and mean number of ripple peaks per ripple event were calculated.

### Statistical analysis

Analysis of the Thio-T aggregation endpoint assay was performed in GraphPad Prism 10.1.2. Different control/tau preparations were compared with a one-way ANOVA, followed by a Tukey’s post-hoc test for multiple pair-wise comparisons.

SWR analysis was performed in MATLAB 2023b (The MathWorks, Inc.) using a generalized linear mixed-effects model (glme; fitglme.m) with normal distribution, restricted maximum pseudo likelihood (REMPL) method for estimating model parameters, and reference coding for dummy variables. The model equations used were:

a. For the pre-incubation experiments: SWR_parameter ∼ Condition (Control or Tau) + (1|SubjectID) + (-1 + Condition|SubjectID)
b. For the acute tau application experiments: SWR_parameter ∼ Condition (pre, tau, and post) + (1|SubjectID) + (1|SliceID) + (-1 + Condition|SubjectID)

where (1|SubjectID) is a subject-specific random intercept that captures each subject’s overall mean response, (-1 + Condition|SubjectID) allows different sensitivity of each subject to each condition that is uncorrelated to the subject-specific intercept, and (1|SliceID) is an additional random intercept for slices (needed only when each slice is exposed to all different conditions (baseline pre-tau, application of tau, and after washout) and hence has repeated measures). We tested for an overall effect of condition (fixed effect) using an ANOVA on the glme. Since there are only 2 conditions in the pre-incubation experiments, no post-hoc test was necessary, while for the acute application experiments pairwise comparisons between conditions were performed using Holm-Bonferroni-corrected post-hoc tests. P-values and F-statistics are reported in two significant figures. Condition-level estimated marginal means (EMMs, model-based condition means adjusted for random effects) and their standard errors (SEs) were derived from the glme.

To calculate the log_10_(fold-change) of SWR parameters during acute application of otau on mouse and human hippocampal slices, all SWR parameters were log_10_-transformed before fitting the glme as in equation b, above. The log_10_(fold-change) relative to baseline/pre-tau corresponds to the estimated marginal means (EMMs) for each fixed-effect contrast (tau or washout/post-tau) from this model. The 95% confidence intervals (CIs) were computed using standard errors derived from the covariance matrix of the fixed effects, multiplied by the critical value of the t-distribution at 2.5% in each tail, with degrees of freedom obtained from the model.

Comparison of pre-tau mouse recordings from acute experiments and control recordings from pre-incubation experiments allowed assessment of the effects of reusing the ACSF in pre-incubation experiments. EMMs and SEs were calculated from a glme with equation:

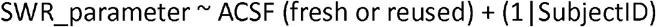

Lastly, to test if between-subject variability differed between mice and humans, mean SWR responses were calculated per subject, and a Monte Carlo permutation test was performed. In each of the 10,000 permutations, species labels were randomly shuffled while the original number of subjects in each species was preserved, and the SD difference between species (SD_mice - SD_humans) was calculated. The two-sided p-value was computed as the proportion of permutations in which the absolute permuted difference was equal to or greater than the observed difference between SDs.

## Results

### Tau oligomers decrease ripple duration in human hippocampal slices

To test how human tau oligomers would affect hippocampal slices, we bath-applied otau-containing ACSF while recording from slices that already had spontaneously generated SWRs. SWRs were analyzed from 2-minute snippets taken before tau application (pre), after 30-minutes of tau-ACSF (tau), and after 30-minutes of washout (post). Because different patients had different underlying conditions and genetic backgrounds, this within-subjects design is ideal. We saw a 12% decrease in ripple duration (*p* = 0.017; Fig. 3A) during otau application, which recovered back to baseline levels after washout in tau-free ACSF (pre = 49 ± 5.3 ms, tau = 43 ± 5.3 ms, post = 51 ± 5.5 ms). The same trend of a 12% decrease was apparent for the number of ripple peaks per ripple event (*p* = 0.024; Fig. 3B; *p*_pre-tau = 0.053; *p*_pre-post = 0.54; *p*_tau-post = 0.050; pre = 13.2 ± 1.5 peaks, tau = 11.6 ± 1.5 peaks, post = 13.8 ± 1.6 peaks), while ripple frequency was unaffected (Fig. 3C), suggesting the decreased ripple duration was due to decreased ripple cycles per event rather than altered ripple frequency. Ripple power (Fig. 3D), amplitude (Fig. 3E), and SW FWHM (Fig. 3F) were unaffected by tau. SW amplitude decreased (*p* = 0.0035; Fig. 3G), with post-hoc comparison finding a significant difference only between pre and post conditions (30% decrease with pre = 47 ± 9.1 μV, tau = 41 ± 10 μV, post = 33 ± 9.1 μV). The rate of SWR occurrence was not changed across conditions (Fig. 3H). Figure 4 shows a representative SWR trace and event from the same slice during pre, tau, and post, with asterisks highlighting detected ripple peaks to demonstrate the decreased ripple duration.

**Figure 2:**
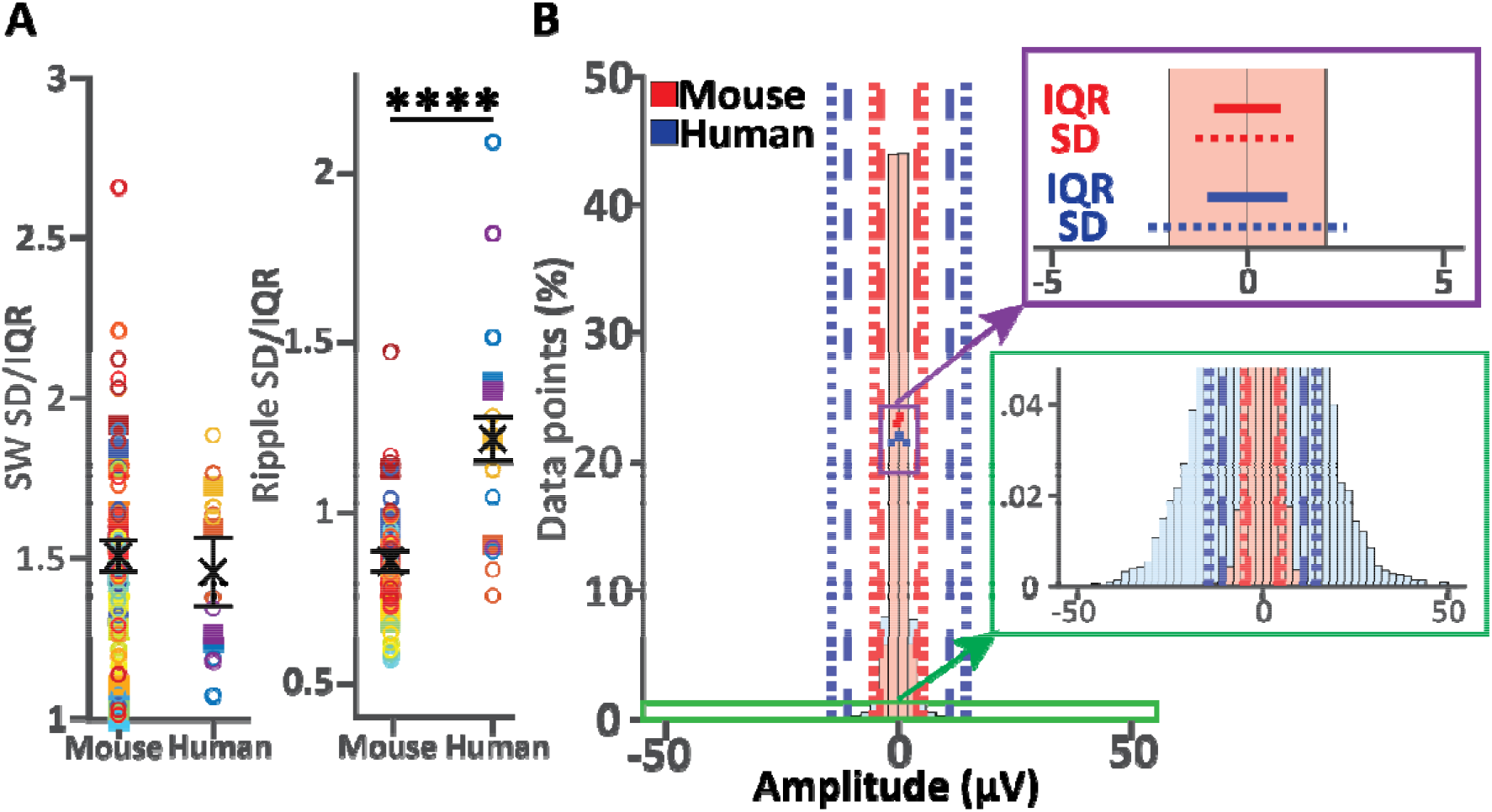
Ripple SD/IQR, but not SW SD/IQR, was higher in human recordings compared to mouse recordings. **(A)** Each graph shows estimated marginal means (EMMs) ± standard errors (SEs) per species from the glme in black. Filled squares represent subject data, while open circles represent slice data, and different colors are data from different subjects. SW SD/IQR (left) was similar between mouse and human recordings (*F*(1,65) = 0.19, *p* = 0.67). Ripple SD/IQR (right) was significantly higher in humans (*F*(1,65) = 25, *p* = 4.1 × 10^-6^). The ratio of the means (1.22/0.86 = 1.42) suggests that ripple amplitude detection thresholds should be 1.42 times larger in human compared to mouse recordings. The overall effect of species was assessed with ANOVA on the glme. N = 55 slices from 23 mice, 12 slices from 4 humans. (**B**) Histogram of the amplitude of every single point of the 2-minute ripple-filtered recording from a representative mouse (red) and human (blue) slice. Vertical dashed lines show the thresholds used for detection of all ripple peaks (mouse = 3 x SD, human = 4.3 x SD) and vertical dotted lines show the thresholds used for the maximum ripple peak criterion (mouse = 4 x SD, human = 5.7 x SD). The purple rectangle insert is a zoom of the IQR (horizontal solid line) and SD (horizontal dotted line) of each slice, to demonstrate that the two slices have the same IQR, suggesting similar baseline noise, but the SD of the human slice is larger, suggesting larger signal, resulting in a larger SD/IQR ratio. We took this as an indication of a higher signal to noise ratio in human slices, and hence that a larger threshold is needed to detect human ripples. The green rectangle insert shows a close-up of the tails of the two slices. These tails represent the ripple events (i.e., the signal), illustrating why the larger threshold used in human slices is appropriate.

**Figure 3:**
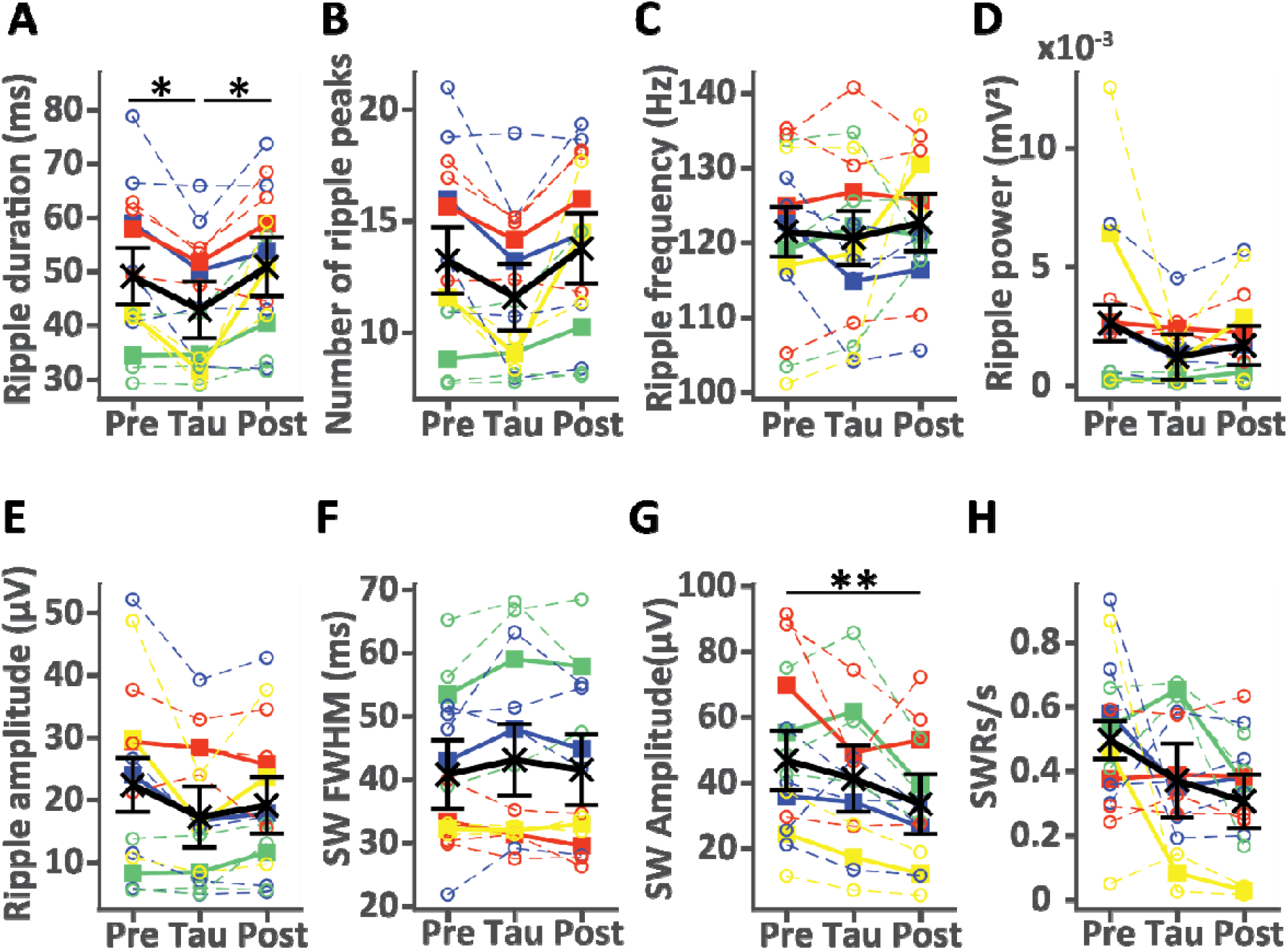
Acute application of otau on human hippocampal slices decreased ripple duration. Each graph shows estimated marginal means (EMMs) ± standard error (SEs) per condition from the generalized linear mixed-effects (glme) model in black. Solid lines with filled squares represent subject data, while dotted lines with open circles represent slice data, and different colors are data from different subjects. Ripple duration (**A**) decreased during tau application and recovered back to pre-tau levels during washout (*F*(2,33) = 4.6, *p* = 0.017). The same trend was seen in the number of ripple peaks per ripple event (**B**; *F*(2,33) = 4.2, *p* = 0.024; *p*_pre-tau = 0.053; *p*_pre-post = 0.54; *p*_tau-post = 0.050). Exposure to otau did not affect ripple frequency (**C**; *F*(2,33) = 0.22, *p* = 0.81), ripple power (**D**; *F*(2,33) = 1.6, *p* = 0.22), ripple amplitude (**E**; *F*(2,33) = 1.8, *p* = 0.19), or SW full-width at half-maximum (**F**; *F*(2,33) = 0.71, *p* = 0.50). SW amplitude (**G**) decreased from pre-tau to post-tau (*F*(2,33) = 6.7, *p* = 0.0035). SWR rate (**H**) was unaffected by tau application (*F*(2,33) = 2.0, *p* = 0.16). The overall effect of condition was assessed with ANOVA on the glme. Pairwise differences were evaluated with Holm–Bonferroni–corrected post-hoc tests with * = *p* < 0.05 and ** = *p* < 0.01. N = 12 slices from 4 patients.

**Figure 4:**
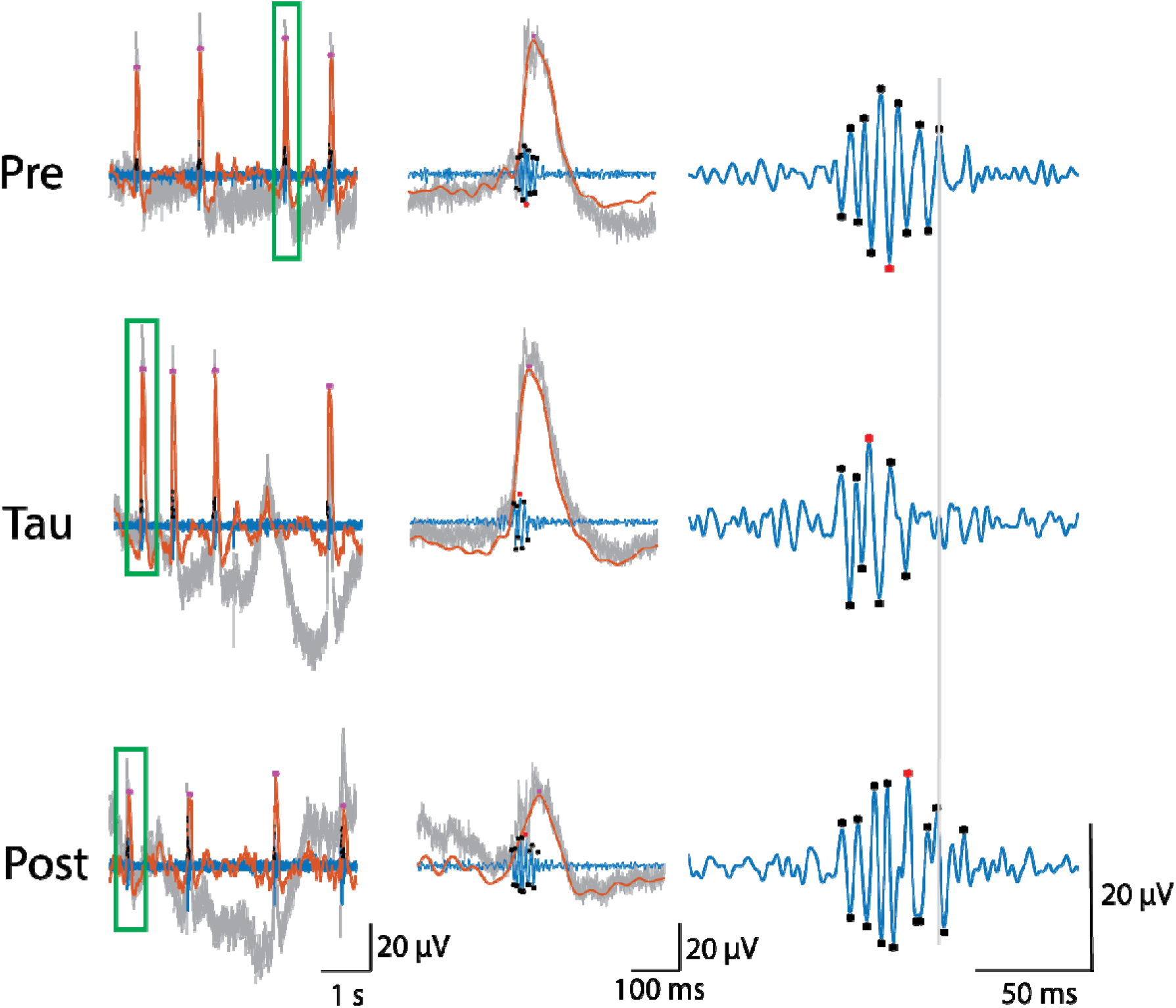
Representative traces and SWR events from the same human slice across the three different conditions of baseline, otau application, and washout. Raw (grey traces), ripple filtered (70–250 Hz, blue traces) and SW filtered (1–25 Hz, orange traces) recordings from baseline (top row, pre-tau), otau application (middle row) and washout (bottom row, post-tau). The left column shows a five second example trace of human SWRs. The green rectangle shows the selected SWR event that is zoomed into in the middle and right columns. Black asterisks mark detected ripple peaks, with the red asterisk marking the largest ripple peak of each event, and magenta asterisks mark detected SW peaks. Ripple asterisks are only positive in the left column because they were detected from the rectified ripple channel, but in the middle and right columns they were moved to match their respective negative peaks. Events in the right column are aligned to the first ripple peak, while the grey vertical line is aligned to the last ripple peak of the baseline condition. Note the decreased ripple duration (time of last peak - time of first peak), as well as the tendency for decreased number of ripple peaks per event, during otau application compared to pre- and post-tau conditions.

### Tau oligomers decrease SWR rate, amplitude, and power in mouse hippocampal slices

When we performed the same experiment in mouse hippocampal slices, ripple duration (Fig. 5A), number of ripple peaks per ripple event (Fig. 5B), and ripple frequency (Fig. 5C) were unaffected by otau. However, ripple power (*p* = 0.0014; Fig. 5D) and amplitude (*p* = 0.0036; Fig. 5E) significantly decreased from baseline after otau application and partially recovered during washout (for power: 23% reduction from pre = 1.3 ± 0.18 × 10^-3^ mV^2^, tau = 1.0 ± 0.18 × 10^-3^ mV^2^, post = 1.1 ± 0.18 × 10^-3^ mV^2^ ; for amplitude: 14% reduction with pre = 14 ± 1.3 μV, tau = 12 ± 1.4 μV, post = 13 ± 1.5 μV). Tau did not change SW FWHM (Fig. 5F) but decreased SW amplitude (*p* = 0.010; Fig. 5G) and SWR rate (*p* = 0.00079; Fig. 5H). Both recovered only partially after washing in tau-free ACSF (for SW amplitude: 18% reduction from pre = 96 ± 14 μV, tau = 79 ± 15 μV, post = 84 ± 16 μV; for SWR rate: 30% reduction from pre = 1.2 ± 0.16 SWRs/s, tau = 0.84 ± 0.16 SWRs/s, post = 1.0 ± 0.16 SWRs/s). To demonstrate these changes, traces and events from a representative slice are shown in Fig. 6. Changes in ripple power are illustrated by the mean power spectral density (PSD) of an example slice across each condition (Fig. 7). Figure 8 summarizes the results of acute application of otau on mouse and human hippocampal slices.

**Figure 5:**
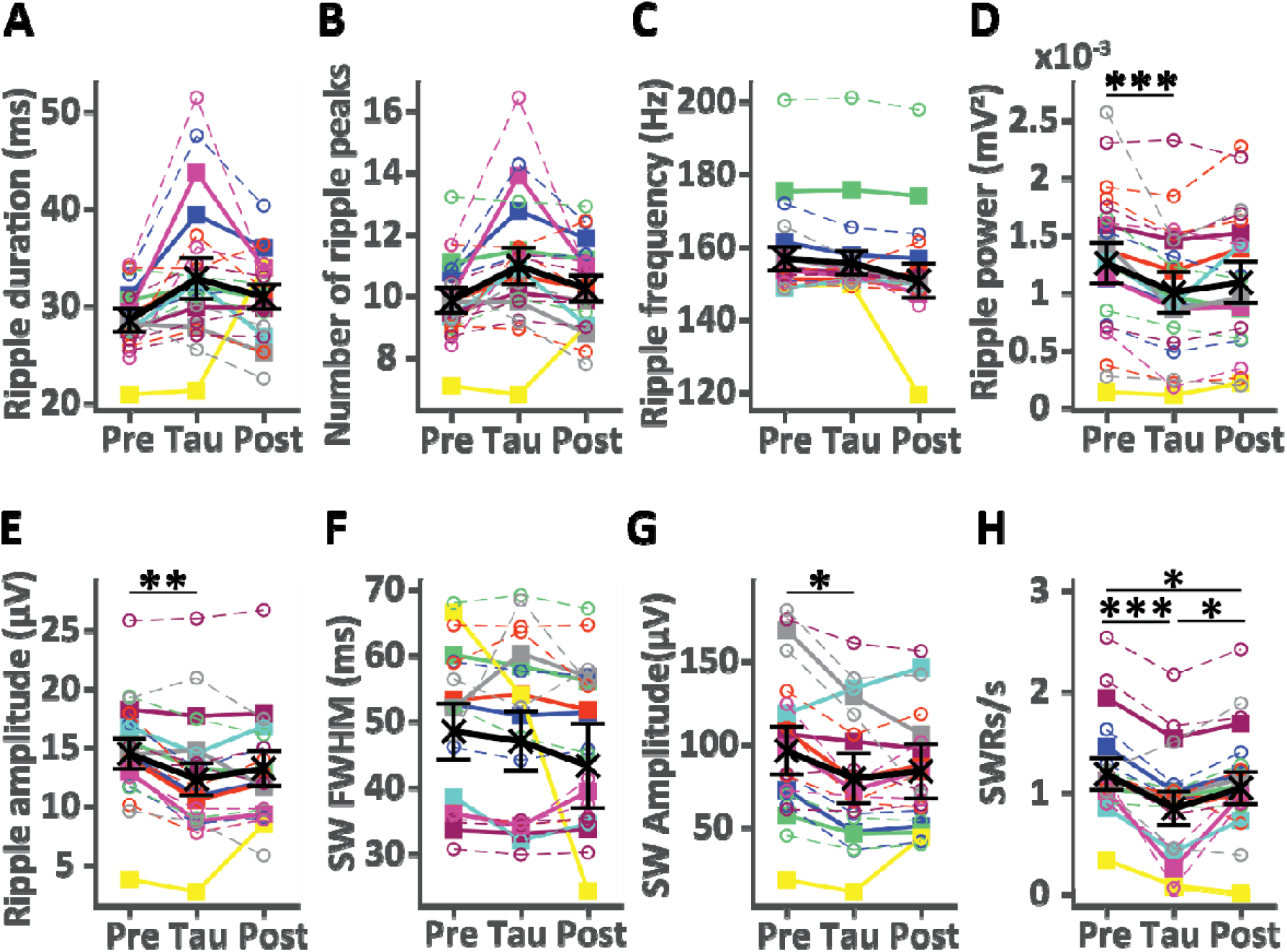
Acute application of otau on mouse hippocampal slices decreased SWR rate, amplitude, and power, but spared duration. Each graph shows EMMs ± SEs per condition from the glme model in black. Solid lines with filled squares represent subject data, while dotted lines with open circles represent slice data, and different colors are data from different subjects. Ripple duration (**A**; *F*(2,45) = 2.5, *p* = 0.095), number of ripple peaks per ripple event (**B**; *F*(2,45) = 2.1, *p* = 0.13), and ripple frequency (**C**; *F*(2,45) = 1.7, *p* = 0.19) were unaffected by exposure to tau. Ripple power (**D**; *F*(2,45) = 7.6, *p* = 0.0014) and ripple amplitude (**E**; *F*(2,45) = 6.4, *p* = 0.0036) decreased during bath application of otau and partially recovered during washout. SW full-width at half-maximum (**F**) did not change across conditions (*F*(2,45) = 0.55, *p* = 0.58). SW amplitude (**G**; *F*(2,45) = 5.1, *p* = 0.010) and SWR rate (**H**; *F*(2,45) = 8.4, *p* = 0.00079) decreased during tau and partially recovered during washout. The overall effect of condition was assessed with ANOVA on the glme. Pairwise differences were evaluated with Holm–Bonferroni–corrected post-hoc tests with * = *p* < 0.05, ** = *p* < 0.01, and *** = *p* < 0.001. N = 16 slices from 8 mice.

**Figure 6:**
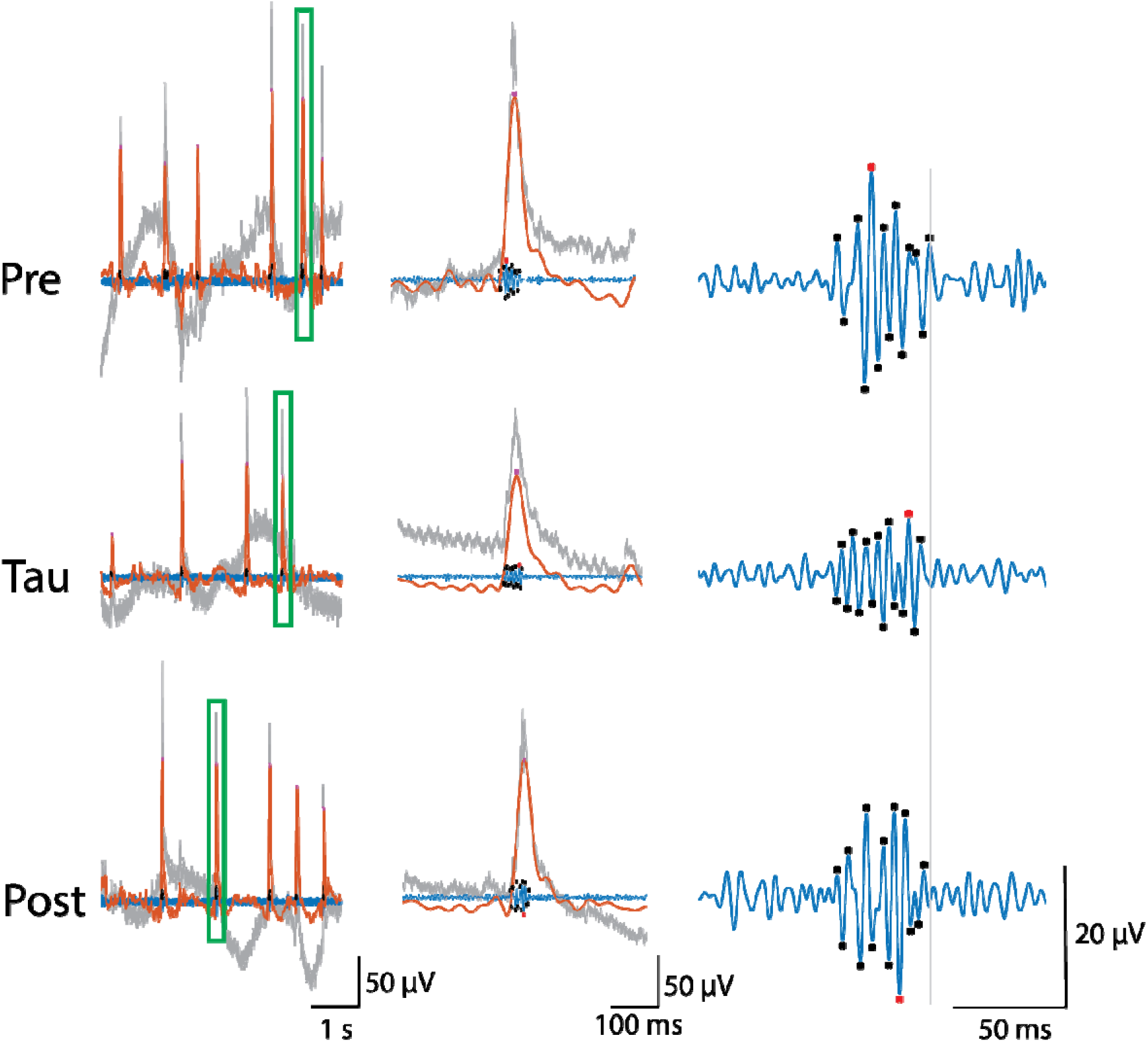
Representative traces and SWR events from the same mouse slice across the three different conditions of baseline, otau application, and washout. Raw (grey traces), ripple filtered (100–250 Hz, blue traces) and SW filtered (1–25 Hz, orange traces) recordings from baseline (top row, pre-tau), otau application (middle row) and washout (bottom row, post-tau). The left column shows a five second trace from each recording to demonstrate the decreased SWR rate during tau. The green rectangle shows the selected SWR event that is zoomed into in the middle column to show the decreased SW amplitude during tau, with further zoom in the right column to show the decreased ripple amplitude of the maximum ripple peak of each event (red asterisks) during tau. The grey vertical line in the right column is aligned to the last ripple peak of the pre-tau event. Magenta asterisks show SWs and black asterisks show ripple peaks detected automatically. Ripple asterisks are only positive in the left column because they were detected from the rectified ripple channel, but in the middle and right columns they were moved to match their respective negative peaks. Note the decreased SWR rate, SW amplitude, and ripple amplitude during otau application compared to baseline and that SWRs partially recovered during washout.

**Figure 7:**
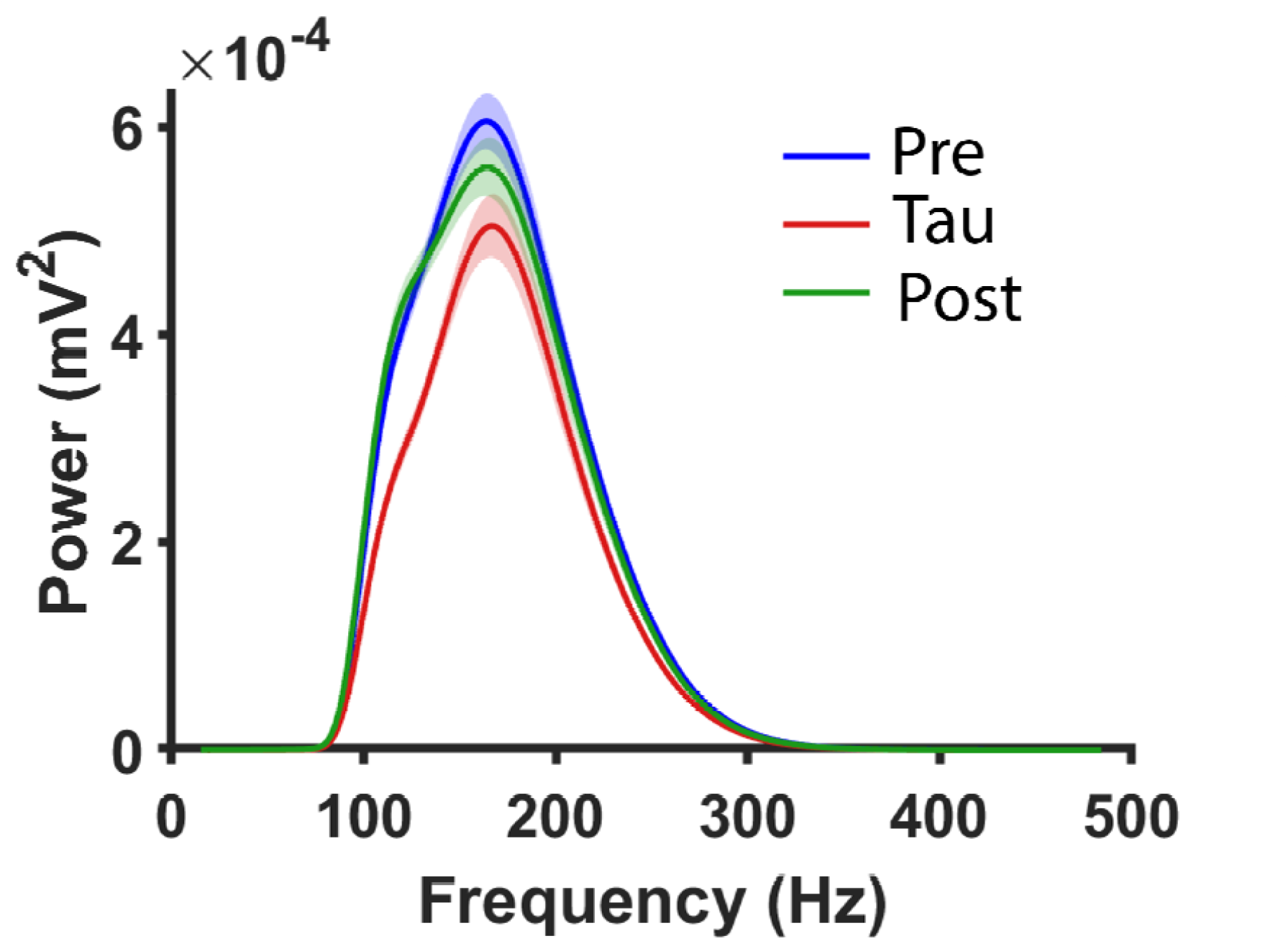
Power spectrum density (PSD) graph demonstrating decreased ripple power during application of tau. The graph shows mean ± SE of the PSD of all events from a single slice across the three conditions. Note the decreased ripple power during otau application that partially recovered after washout (post).

**Figure 8:**
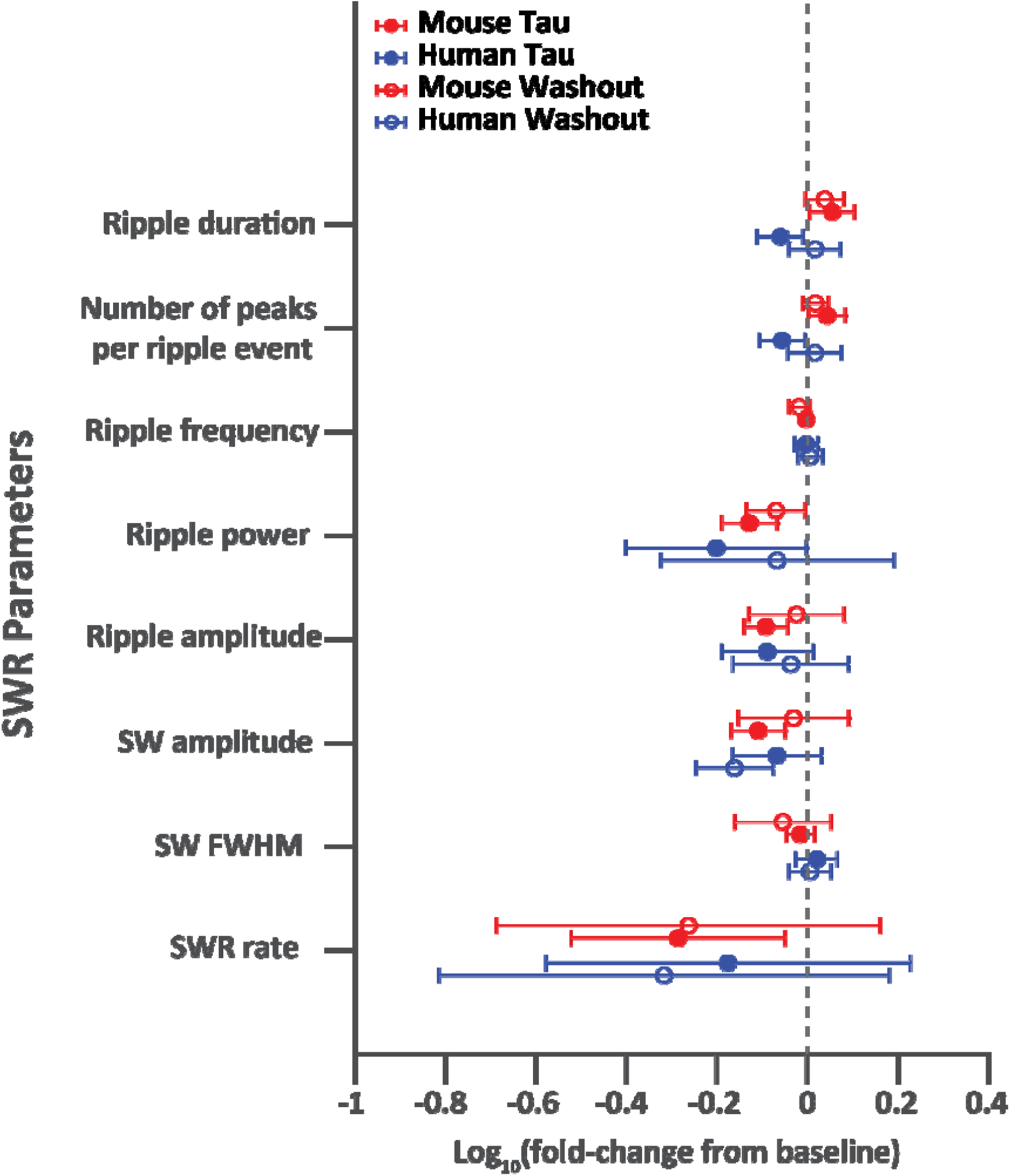
Result summary of acute otau application on mouse and human hippocampal slices. The graph shows log_10_ fold-change ± 95% confidence interval (CI) of SWR parameters (y axis) from baseline (pre) to tau (filled circles) and from baseline (pre) to washout (post; empty circles) for human (blue) and mouse (red) hippocampal slices. Vertical line at 0 represents baseline.

### Pre-incubation of mouse slices with tau oligomers decreases SWR rate

It is possible that if slices were exposed to tau immediately after slicing, SWR generation would be blocked. Therefore, while not as powerful as a within-subjects design, we decided to incubate mouse hippocampal slices in either tau-ACSF or tau-free ACSF directly after slicing. Slices pre-incubated with otau still generated SWRs that could be recorded 2.5–5.5 hours later, albeit at a decreased incidence rate (*p* = 0.029; 37% reduction; control = 0.89 ± 0.16 SWRs/s, tau = 0.56 ± 0.15 SWRs/s), while leaving other SWR parameters unchanged (Fig. 9). The decreased SWR rate in slices pre-incubated with otau is also seen in representative example slices in Fig. 10.

**Figure 9:**
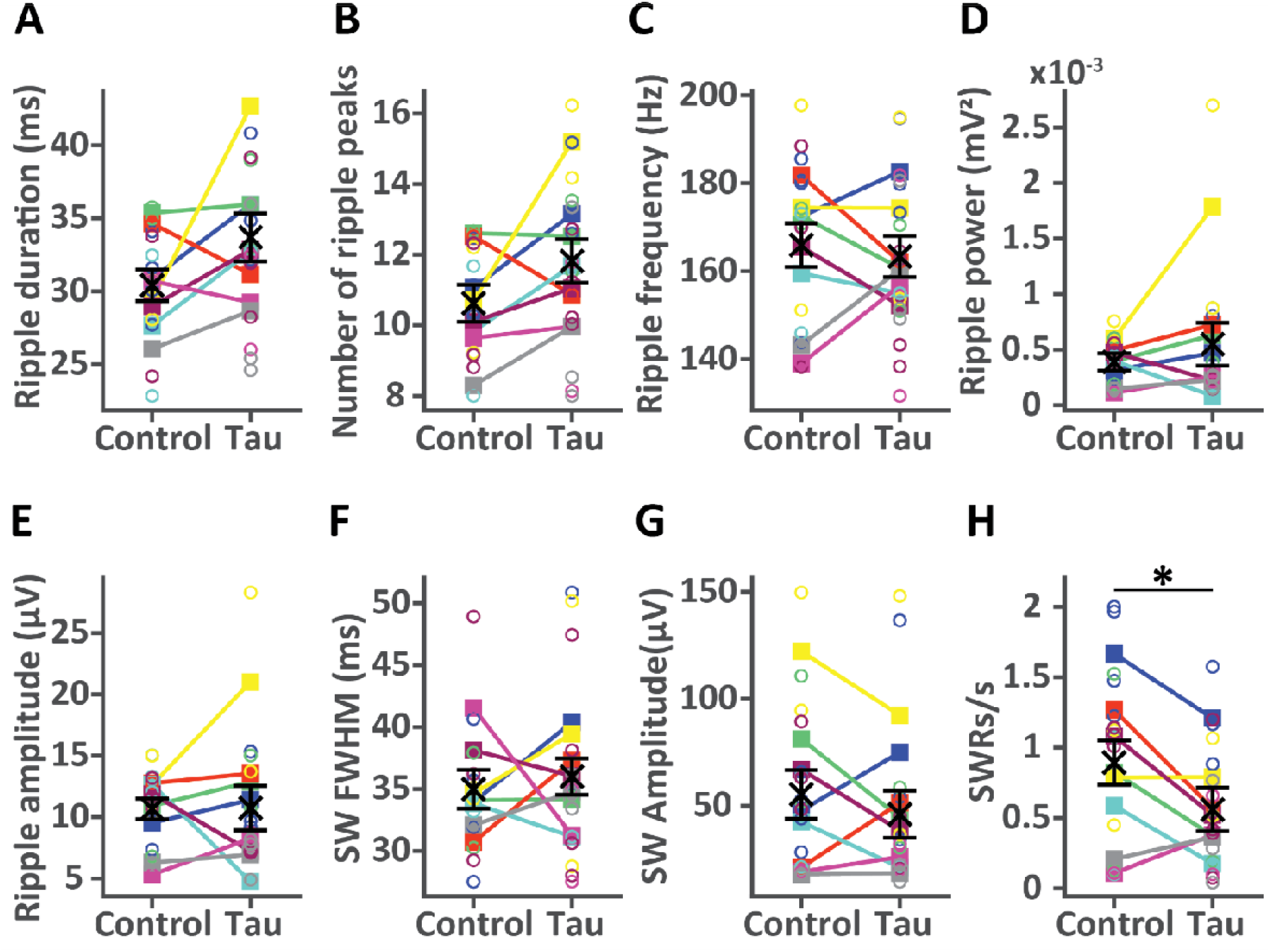
Pre-incubation of mouse hippocampal slices with otau only impaired SWR rate. Each graph shows EMMs ± SEs per condition from the glme in black. Solid lines with filled squares represent subject data, while open circles represent slice data, and different colors are data from different subjects. Ripple duration (**A**; *F*(1,32) = 2.9, *p* = 0.10), number of ripple peaks per ripple event (**B**; *F*(1,32) = 2.8, *p* = 0.10), ripple frequency (**C**; *F*(1,32) = 0.16, *p* = 0.69), ripple power (**D**; *F*(1,32) = 0.59, *p* = 0.45), ripple amplitude (**E**; *F*(1,32) = 0.0013, *p* = 0.97), SW full-width at half-maximum (**F**; *F*(1,32) = 0.22, *p* = 0.64), and SW amplitude (**G**; *F*(1,32) = 0.74, *p* = 0.40) were unaffected by exposure to tau. SWR rate (**H**) was reduced in slices incubated with otau ACSF compared to control ACSF (*F*(1,32) = 5.2, *p* = 0.029). The overall effect of condition was assessed with ANOVA on the glme with * = *p* < 0.05. N = 16 control and 18 tau slices from 8 mice.

**Figure 10:**
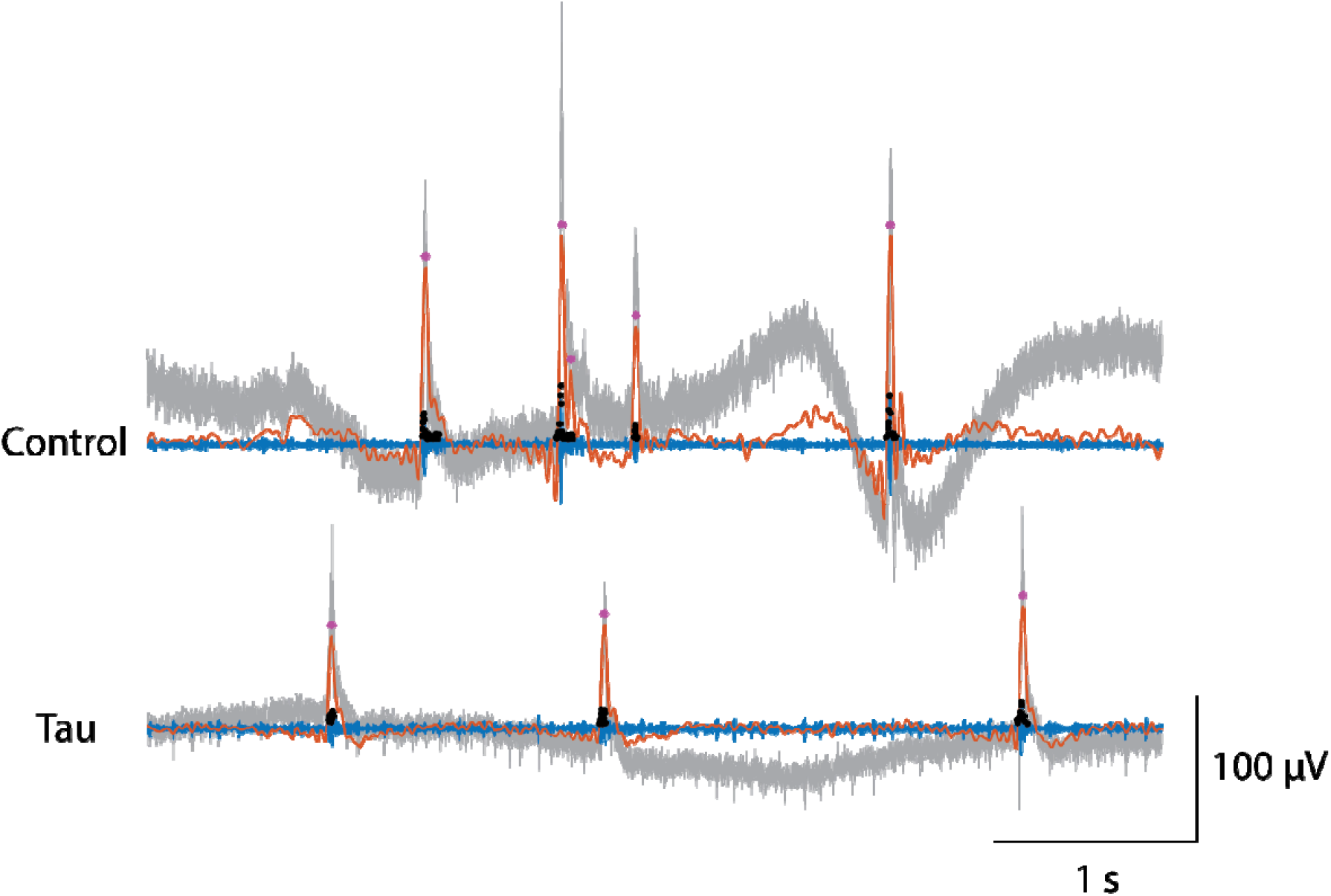
Representative traces demonstrating decreased SWR rate in slices pre-incubated with otau. Raw (grey traces), ripple filtered (100–250 Hz, blue traces) and SW filtered (1–25 Hz, orange traces) recordings from a slice pre-incubated in control ACSF (top) and a slice pre-incubated in ACSF containing otau (bottom) from the same mouse, to show the decreased SWR rate in slices pre-incubated with tau oligomers. Magenta asterisks show detected SWs and black asterisks show detected ripple peaks (only positive because they were detected from the rectified ripple channel).

### A tau preparation lacking β-sheets had no effect on SWRs

We also tested oligomeric tau prepared with hydrogen peroxide (otau-H_2_O_2_), as tau oligomers prepared with this method were shown to decrease LTP in mouse hippocampal slices (16,17). However, in our hands, this preparation lacked β-sheets, as shown by Thio-T assay (Fig. 1). Thus, it served as a control of non-aggregated tau. Neither acute application of otau-H_2_O_2_ for 30 minutes (Fig. 11), nor pre-incubation of slices with otau-H_2_O_2_ directly after slicing (Fig. 12) had any effect on SWRs.

**Figure 11:**
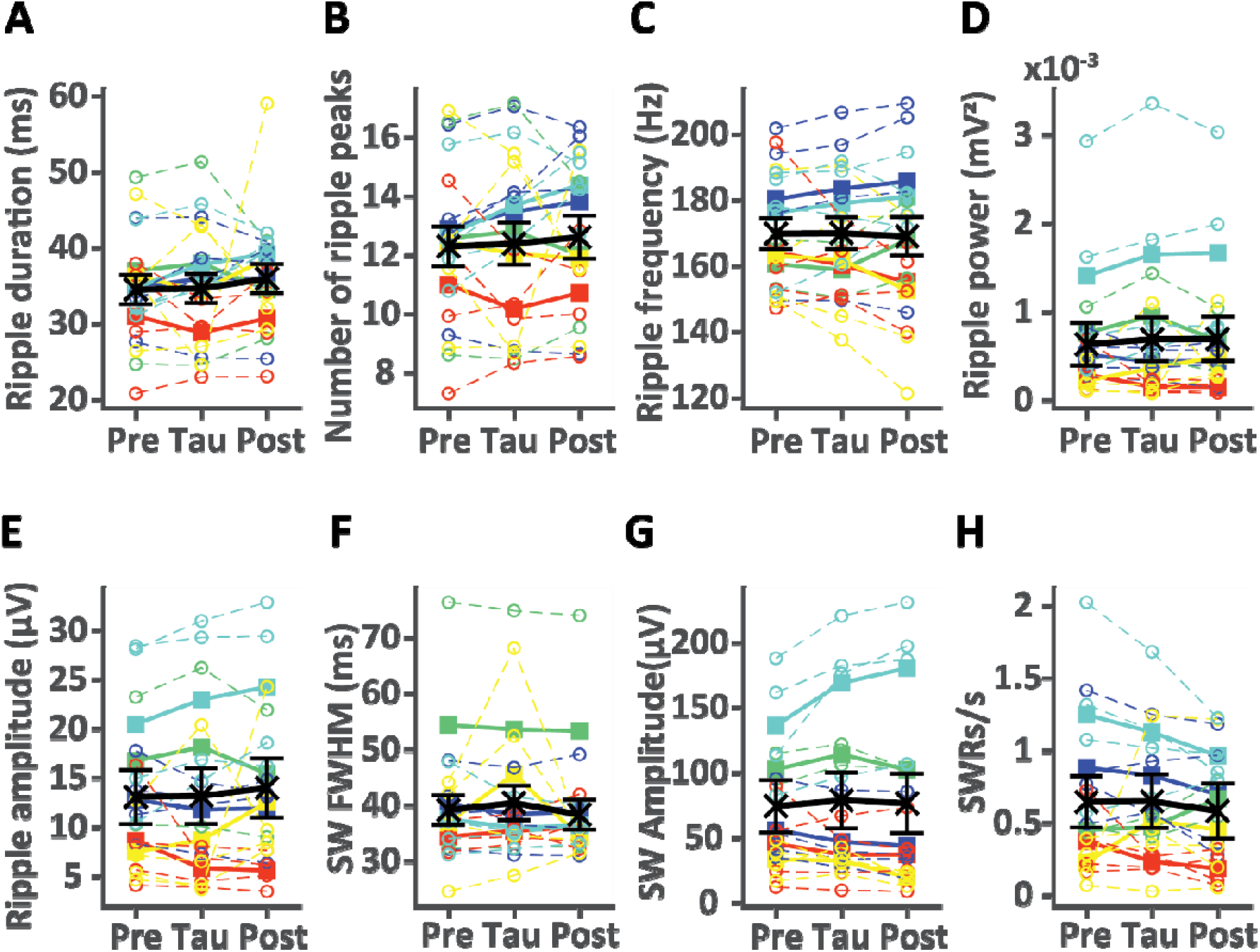
Acute application of otau-H_2_O_2_ on mouse hippocampal slices did not affect SWRs. Each graph shows EMMs ± SEs per condition from the glme in black. Solid lines with filled squares represent subject data, while dotted lines with open circles represent slice data, and different colors are data from different subjects. Ripple duration (**A**; *F*(2,51) = 0.37, *p* = 0.70), number of ripple peaks per ripple event (**B**; *F*(2,51) = 0.17, *p* = 0.85), ripple frequency (**C**; *F*(2,51) = 0.041, *p* = 0.96), ripple power (**D**; *F*(2,51) = 0.22, *p* = 0.80), ripple amplitude (**E**; *F*(2,51) = 0.22, *p* = 0.80), SW full-width at half-maximum (**F**; *F*(2,51) = 0.38, *p* = 0.69), SW amplitude (**G**; *F*(2,51) = 0.45, *p* = 0.64) and SWR rate (**H**; *F*(2,51) = 0.49, *p* = 0.62) were unaffected by exposure to tau. The overall effect of condition was assessed with ANOVA on the glme. N = 18 slices from 5 mice.

**Figure 12:**
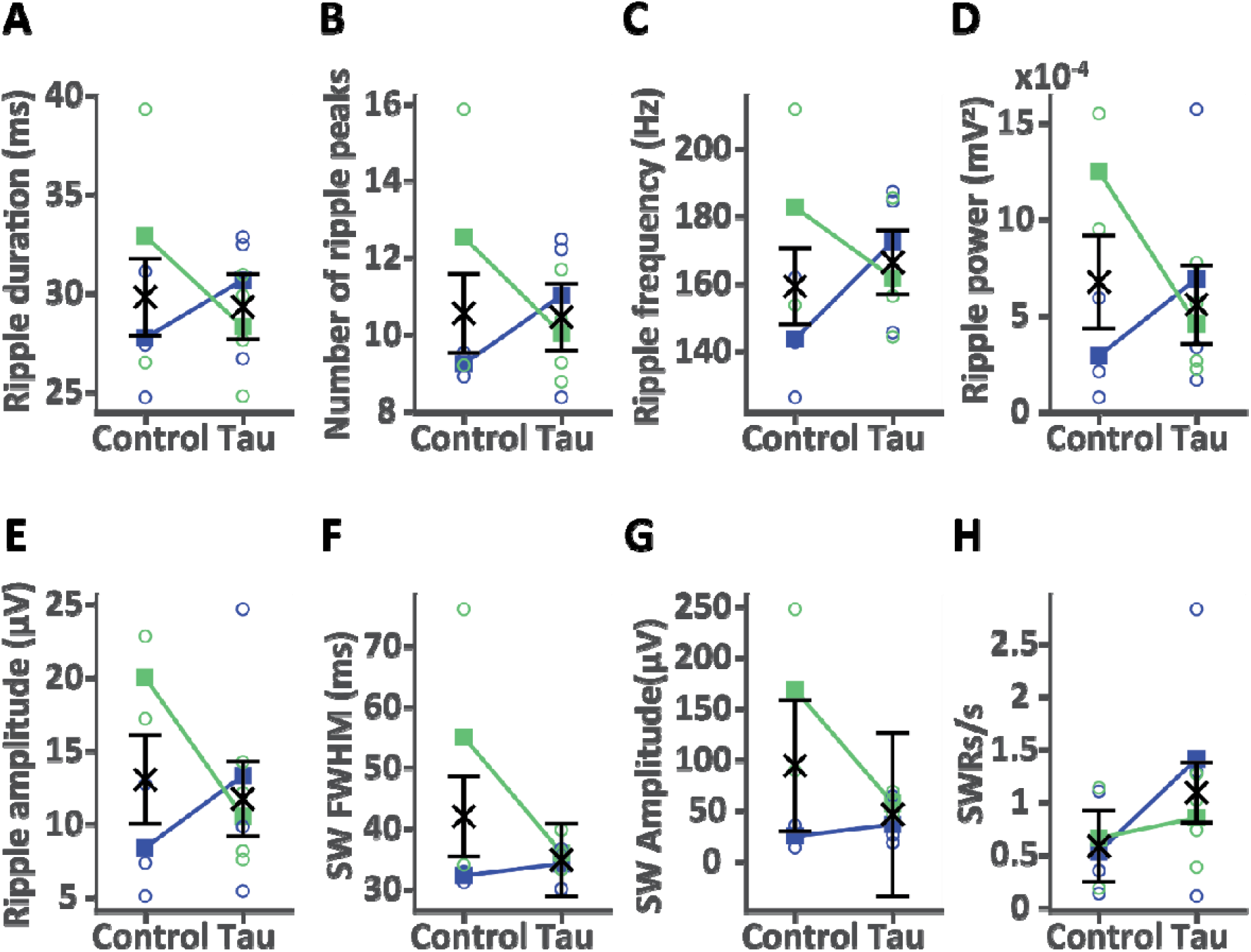
Pre-incubation of mouse hippocampal slices with otau-H_2_O_2_ did not affect SWRs. Each graph shows EMMs ± SEs per condition from the glme in black. Solid lines with filled squares represent subject data, while open circles represent slice data, and different colors are data from different subjects. Ripple duration (**A**; *F*(1,10) = 0.036, *p* = 0.85), number of ripple peaks per ripple event (**B**; *F*(1,10) = 0.0054, *p* = 0.94), ripple frequency (**C**; *F*(1,10) = 0.23, *p* = 0.64), ripple power (**D**; *F*(1,10) = 0.14, *p* = 0.72), ripple amplitude (**E**; *F*(1,10) = 0.11, *p* = 0.75), SW full-width at half-maximum (**F**; *F*(1,10) = 1.1, *p* = 0.33), SW amplitude (**G**; *F*(1,10) = 0.76, *p* = 0.40), and SWR rate (**H**; *F*(1,10) = 1.3, *p* = 0.28) were unaffected by exposure to tau. The overall effect of condition was assessed with ANOVA on the glme. N = 5 control and 7 tau slices from 2 mice.

### Reusing the ACSF impaired ripple power

In pre-incubation experiments, where oligomeric tau application was more prolonged, control- and tau-ACSF solutions were reused to conserve tau. Reusing the ACSF might harm the slices due to exposure to toxic metabolites; and this could then affect how tau changes SWRs. We therefore tested if control mouse SWRs were different in pre-incubation experiments, where the ACSF was reused, compared to pre-tau mouse recordings from acute experiments, where only fresh ACSF was used. We saw a decrease in ripple power in slices where the ACSF was reused (*p* = 0.022; 52% decrease with acute = 0.95 ± 0.13 × 10^-3^ mV^2^, pre-incubation = 0.46 ± 0.16 × 10^-3^ mV^2^), with no significant changes in other SWR parameters (Fig. 13). However, a trend for decreased SW FWHM with ACSF reuse was found (*p* = 0.053; 16% decrease with acute = 44 ± 2.5 ms, pre-incubation = 37 ± 3.0 ms).

**Figure 13:**
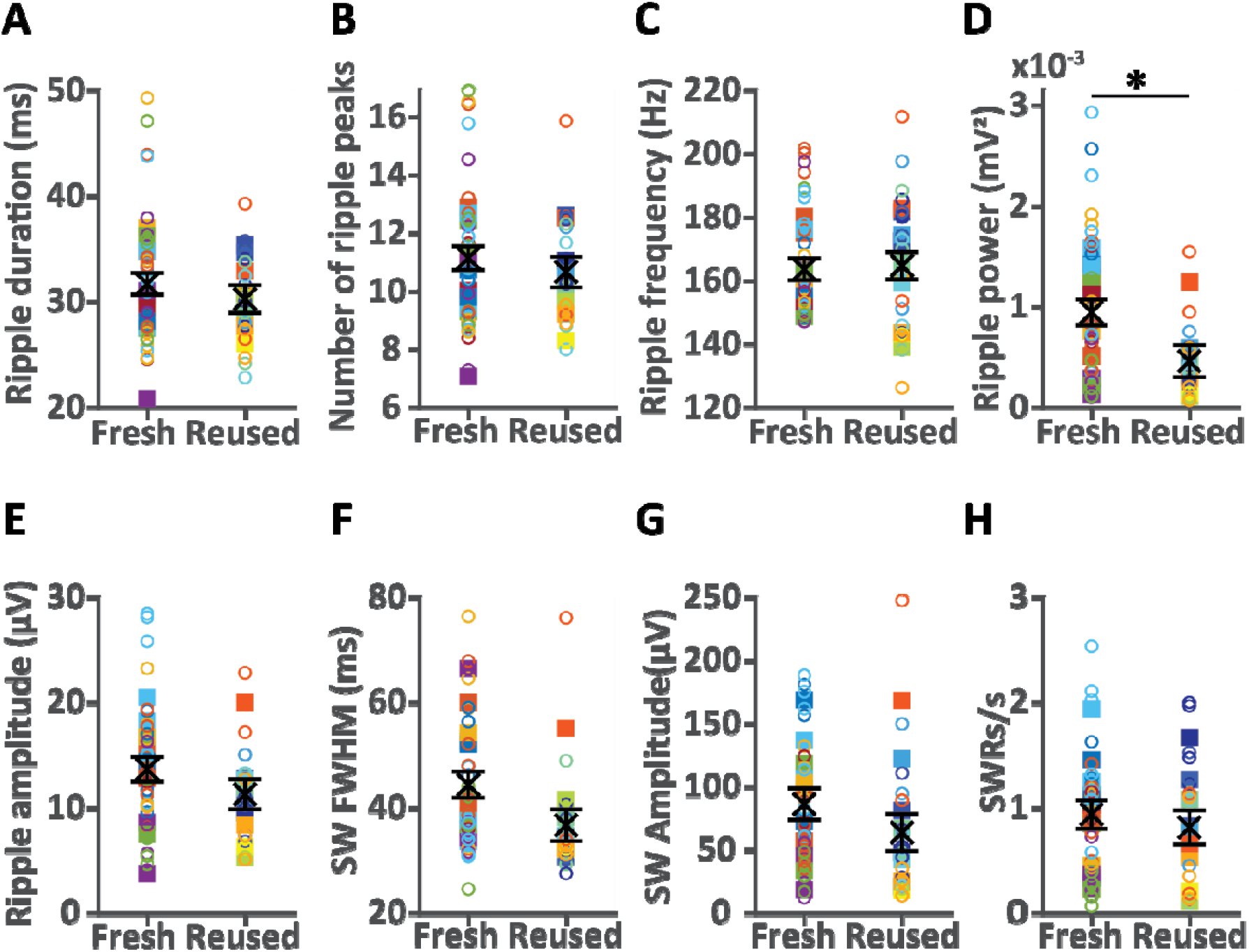
Ripple power was decreased in mouse slices where tau-free ACSF was reused. Each graph shows EMMs ± SEs per condition from the glme in black. Filled squares represent subject data, while open circles represent slice data, and different colors are data from different subjects. Reusing the ACSF during pre-incubation experiments did not affect ripple duration (**A**; *F*(1,53) = 0.75, *p* = 0.39), number of peaks per ripple event (**B**; *F*(1,53) = 0.51, *p* = 0.48), or ripple frequency (**C**; *F*(1,53) = 0.043, *p* = 0.84), but significantly decreased ripple power (**D**; *F*(1,53) = 5.6, *p* = 0.022) compared to baseline recordings from acute tau application experiments with fresh ACSF. Ripple amplitude (**E**; *F*(1,53) = 1.6, *p* = 0.21), SW full-width at half-maximum (**F**; *F*(1,53) = 3.9, *p* = 0.053), SW amplitude (**G**; *F*(1,53) = 1.4, *p* = 0.25), and SWR rate (**H**; *F*(1,53) = 0.34, *p* = 0.56) were not significantly different between experiments. The overall effect of condition was assessed with ANOVA on the glme with * = *p* < 0.05. N acute = 34 slices from 13 mice, N pre-incubation = 21 slices from 10 mice.

### Variability in most SWR parameters between mouse and human subjects was similar

In order to compare variability between mouse and human subjects in each analyzed SWR parameter, the mean response per subject was computed and a permutation test performed on the SD difference (SD_mice - SD_humans). Variability in the number of peaks per ripple event was larger in humans than in mice (SD_mice = 1.58 peaks, SD_humans = 3.44 peaks, *p* = 0.035; Fig. 14B), with the same trend seen in ripple duration (SD_mice = 3.80 ms, SD_humans = 12.1 ms, *p* = 0.060; Fig. 14A). Ripple amplitude was also more variable in humans than in mice (SD_mice = 4.48 μV, SD_humans = 10.1 μV, *p* = 0.035; Fig. 14E), while SWR rate was more variable in mice than humans (SD_mice = 0.48 SWR/s, SD_humans = 0.089 SWR/s, *p* = 0.026; Fig. 14H). The larger variability between mice in SWR rate could be due to combining mouse data from both fresh and reused ACSF, or from both male and female mice (compared with data from only male human subjects). We therefore confirmed that the larger variability in SWR rate in mice persisted when only data from acute experiments (with fresh ACSF) were used (SD_mice = 0.49 SWR/s, SD_diff = 0.40 SWRs/s, *p* = 0.033), as well as when only data from male mice from acute experiments were considered (SD_mice = 0.40 SWR/s, SD_diff = 0.31 SWRs/s, *p* = 0.012). There was no difference in variability between humans and mice in terms of ripple frequency (Fig. 14C), ripple power (Fig. 14D), SW FWHM (Fig. 14F), or SW amplitude (Fig. 14G).

**Figure 14:**
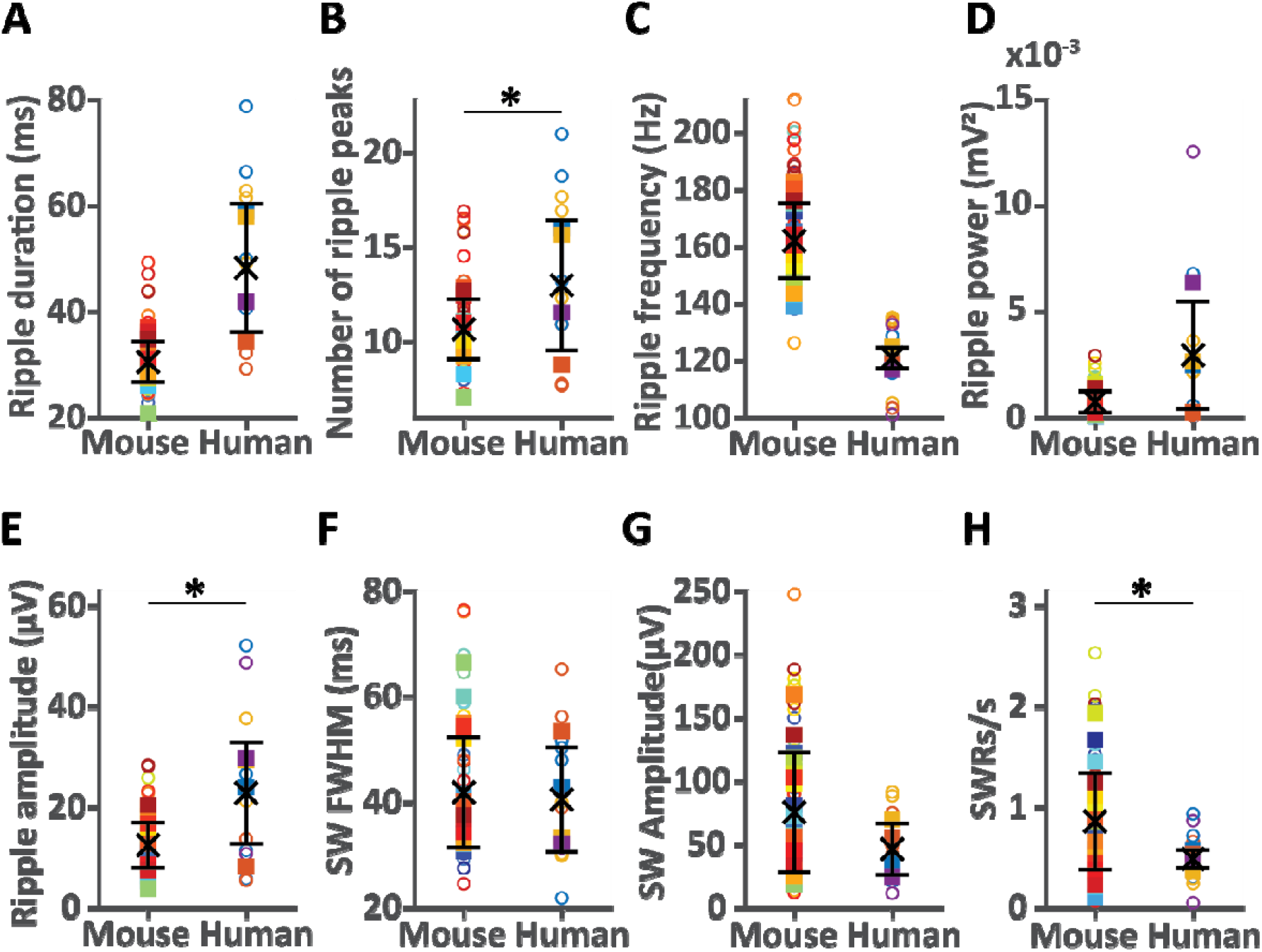
Comparison of SWR variability in mouse and human subjects. Each graph shows mean ± SD calculated from subject-level means in black. Filled squares represent subject data, while open circles represent slice data, and different colors are data from different subjects. Ripple duration (**A**) showed a trend for a larger SD in human compared to mouse subjects, but was not significantly different (SD_diff = -8.3 ms, *p* = 0.060). The number of peaks per ripple event (**B**) was more variable in humans than in mice (SD_diff = -1.9 peaks, *p* = 0.035). The variability in ripple frequency (**C**) was similar between mice and humans (SD_diff = 9.5 Hz, *p* = 0.25) as was ripple power (**D**; SD_diff = -2.0 × 10^-3^ mV^2^, *p* = 0.11). Ripple amplitude (**E**) was more variable in humans than in mice (SD_diff = -5.6 μV, *p* = 0.035). Between-subject variability was similar in mice and humans for SW full-width at half-maximum (**F**; SD_diff = 0.55 ms, *p* = 0.87) and SW amplitude (**G**; SD_diff = 27 μV, p = 0.15). SWR rate (**H**) was more variable in mice than in humans (SD_diff = 0.39 SWRs/s, *p* = 0.026). Statistical differences between the SDs of the two species were assessed with a permutation test with * = *p* < 0.05. N = 23 mouse and 4 human subjects.

## Discussion

In order to study acute effects of tau oligomers on SWRs in the human hippocampus, we bath-applied oligomeric tau on human hippocampal slices that had already generated spontaneous SWRs after incubation in tau-free ACSF. We saw that after 30 minutes of exposure to otau, SWR rate, amplitude, and power were already reduced, similar to what is observed in rTg4510 (tauP301L) mice (10,11). In these mice, SWR power is reduced at 2–4 months (10), when tau oligomers are present, but tangles have not yet formed in the hippocampus (30). Tau oligomers occur prior to neuropathological tau aggregation (12), and their active or passive release may contribute to early synaptic impairments (13). The acute effect of tau oligomers on SWRs that we observe in both mouse and human slices, may therefore recapitulate early stages of Alzheimer’s disease and tauopathies. This acute effect is in agreement with in vivo studies where intrahippocampal injections of tau oligomers in wild-type mice led to neurodegeneration and memory impairment only 24 h later (15).

In human slices, we saw a decrease in ripple duration during tau application instead of a change in SWR rate or amplitude. This highlights a conserved effect of tau oligomers in impairing SWRs in both species after only 30 minutes, as well as differences in the SWR parameters that are affected. One explanation is genuine species differences. For example, denser CA3-CA3 synaptic connectivity and less reliable synaptic transmission in mouse compared to human hippocampal networks (31) could make SWR generation (SWR rate) and the synchrony or number of participating pyramidal cells (SWR amplitude) more susceptible to perturbations in mice. Another possible reason for this difference is the potentially altered circuitry of the human hippocampi from resections that we used in this study. The selective pruning of inhibitory synapses (32) and decreased excitatory drive in GABAergic basket cells (33) seen in epileptic tissue can shift the excitation/inhibition balance towards increased excitation, which directly affects SWRs (34–36). Indeed, epileptic hippocampi (from mice) were reported to have decreased SWR rate and increased SW amplitude, ripple frequency, and number of cycles per ripple event (37,38). This could influence which SWR parameters are affected by tau. Lastly, epileptic events such as pathological ripples and interictal epileptiform discharges share overlapping network mechanisms and parameter characteristics with SWRs. Physiological and pathological events exist in a continuum, making reliable identification and exclusion of abnormal activity challenging (39,40). Indeed, their separation has been the topic of many papers. Some separate them based on their spatiotemporal patterns (SWRs are not present in the dentate gyrus, consist of ≥ 3 cycles and have a narrowband frequency domain at 100–250 Hz, while interictal epileptiform discharges have a broadband between 20–500 Hz (41)), others define only fast ripples, oscillations > 200 Hz, as pathological (42,43), while some consider all oscillations produced by the epileptic hippocampal network as pathogenic (44). Importantly, a previous study recorded SWRs from mouse slices before induction of epileptic activity using high potassium, 4-Aminopyridine (4-AP), zero magnesium, or gabazine. SWRs disappeared, and after a transitory phase, interictal-like events appeared, which had much larger amplitude and duration compared to SWRs. Interestingly, they also report a negative correlation between SWRs and interictal events, i.e., slices with large SWRs produced either small or no interictal events while slices with small, infrequent, or no SWRs produced large interictal events (45). Here, we did not attempt to separate detected events into physiological and pathological, so results should be interpreted with caution. However, the SWR parameter values we observed in human slices appear broadly consistent with prior reports of SWRs and we saw no clear evidence to suggest these events are pathological. In any case, the effect of tau on SWRs in human slices reflects an effect on events in the SWR frequency range, which could be pathological, physiological, or both.

We found a significant decrease in SW amplitude from baseline to washout in human tissue. The amplitude progressively decreased linearly over time (pre > tau > post), which could point to either an irreversible effect of tau oligomers that increases as time passes (e.g., due to tau being absorbed into neurons and inducing mitochondrial damage and cytotoxicity (46) and seeding aggregation of endogenous tau; (47)) or to deterioration of the human tissue. The latter could be a result of longer transportation times of the human tissue from the operating room to sectioning and incubation compared to mice. In agreement with this hypothesis, only one out of six human tissue samples we received from Hamburg had SWRs compared to three out of four from Berlin (shorter transportation time). Nevertheless, the effect of tau on ripple duration is fully reversed after washout, so it cannot be explained by mere tissue degradation. Similarly, partial recovery after washout was observed in all SWR parameters altered during otau application in mouse slices. We therefore focused on ripple duration regarding results from human slices.

Surprisingly, only SWR rate was decreased in mouse slices pre-incubated with otau. One possible explanation is that reusing the same 100⍰ml of ACSF, a step necessary to decrease the amount of tau oligomers needed for this longer experiment, could lower the effective tau concentration of the solution. Reduced bioavailability of tau oligomers can occur through their degradation or further aggregation, both of which can happen over time in ACSF. Oligomers and aggregates could also stick to the equipment, further reducing the effective tau oligomer concentration, leading to a less profound SWR impairment compared to acute experiments. It is also possible that since ripple power was already decreased in slices where the ACSF was reused, pre-incubation with tau caused no additional reduction. However, differences in SWR amplitude were not significant between acute and pre-incubated controlled slices, so the differential effect of tau in pre-incubated versus acute conditions is not fully explained by this difference. Reusing the ACSF can lead to accumulation of ions, metabolites, or waste products, all of which could account for the decreased ripple power seen. Given that the effect of tau was less dramatic in pre-incubation compared to acute mouse experiments and considering other technical limitations (e.g., need for larger sample sizes in a between-subjects design study, less availability of human tissue, necessity to prepare tau-ACSF before confirming that slices have SWRs in pre-incubation experiments), we did not test pre-incubation of human slices with tau.

In this study, we also tested acute application of otau prepared using H_2_O_2_ on mouse slices, a preparation that has been shown to diminish LTP after 20 minutes (17). In our hands, this preparation lacked tau β-sheets as shown by the Thioflavin-T aggregation assay, and also lacked any effect on SWR parameters. These results point to a necessary role of this secondary protein structure for SWRs to be impaired.

Since SWR characteristics differ between primates and rodents (1,23–27) and hence different criteria were used for SWR detection in each species (see Methods), we cannot directly compare mean SWR values between mouse and human hippocampal slices. However, comparison of between-subject variability showed that ripple amplitude and number of peaks per event were more diverse in humans than in mice. This is expected given that human patients were more heterogenous in terms of age, genetic background, and pathology. On the other hand, SWR rate was more diverse in mouse subjects, an effect that remained when mouse data with reused ACSF and from female mice were excluded. One possible reason is the decreased SWR rate observed in epileptic tissue (37,38), resulting in a lower dynamic range of SWR rate in human slices. Alternatively, the lower signal to noise ratio seen in mice (where noisier signal leads to higher variability) and the lower threshold for ripple detection used because of it, could both lead to capturing a broader spectrum of events in mice than in humans.

To conclude, we have shown that short-term application of tau oligomers impaired SWRs in human and mouse hippocampal slices and this depended on tau β-sheet containing oligomers. We propose that this method can be further utilized to study SWR changes after acute exposure to different tau and/or Aβ species. Furthermore, potential therapeutic agents could be applied to slices with impaired SWRs after exposure to such toxic aggregates to test for rescue effects. While this could be tested in vivo, with transgenic mouse models or with intracranial injections of pathogenic species, the in vitro method described here allows for faster, higher throughput results, as well as for comparison of effects on mouse versus human SWRs.

## Data availability

Data analyzed in this paper are available from the corresponding author upon request.

## Acknowledgements

We gratefully acknowledge the Institute of Neurophysiology at the Charité University Medicine Berlin and the involved scientists for providing expertise and equipment, as well as for the collection and preparation of viable human brain tissue for experimental studies. This work was supported by funding from the Studienstiftung des Deutschen Volkes and the IMPRS Neuroscience program of the University of Goettingen to C.V., and from the GO-Bio BMBF, BrightFocus Foundation, and Charité 3^R^| Replace – Reduce – Refine, to C.D. We thank Dr. Nikolaus Maier and Dr. Daniel Parthier for their valuable input on SWR analysis. We also thank Johanna Hintze for her outstanding support with collection and slicing of the human hippocampal tissue.

## Author contributions

C.V. and C.D. conceived and designed the study. C.V. collected and analyzed the data. A.C.O.V.A. assisted with data collection for the acute otau application experiment. J.H. and R.S. prepared all tau oligomer mixtures and performed the Thioflavin-T aggregation endpoint assays. P.F. coordinated the human tissue project. J.O. and T.S. are the neurosurgeons that resected the tissue from the human patients. C.V. wrote the manuscript. All authors edited and reviewed the manuscript. C.D. and S.W. provided funding and materials.

## Declaration of interests

The authors declare no competing interests.

## Notes

### Competing Interest Statement

The authors have declared no competing interest.

